# Spt6 directly interacts with Cdc73 and is required for Paf1C recruitment to active genes

**DOI:** 10.1101/2022.05.04.490663

**Authors:** Mitchell A. Ellison, Matthew S. Blacksmith, Sanchirmaa Namjilsuren, Margaret K. Shirra, Rachel A. Schusteff, Eleanor M. Kerr, Fei Fang, Yufei Xiang, Yi Shi, Karen M. Arndt

## Abstract

Paf1C is a conserved transcription elongation factor that regulates transcription elongation efficiency, facilitates co-transcriptional histone modifications, and impacts molecular processes linked to RNA synthesis, such as polyA site selection. Coupling of the activities of Paf1C to transcription elongation requires its association with RNA polymerase II (Pol II). Mutational studies in yeast identified Paf1C subunits Cdc73 and Rtf1 as important mediators of Paf1C recruitment to Pol II on active genes. While the interaction between Rtf1 and the general elongation factor Spt5 is relatively well-understood, the interactions involving Cdc73 remain to be elucidated. Using an *in vivo* site-specific protein cross-linking strategy, we identified direct interactions between Cdc73 and two components of the elongation complex, the elongation factor Spt6 and the largest subunit of Pol II. Through *in vitro* protein binding assays and crosslinking/mass spectrometry, we show that Cdc73 and Spt6 can interact in the absence of additional factors and propose a binding interface. Rapid depletion of Spt6 dissociated Paf1 from chromatin and altered patterns of Paf1C-dependent histone modifications genome-wide. These results reveal previously unrecognized interactions between Cdc73 and the Pol II elongation complex and identify Spt6 as a key factor contributing to Paf1C recruitment to active genes in *Saccharomyces cerevisiae*.

## INTRODUCTION

Eukaryotic cells deploy a large number of regulatory proteins to establish and maintain a chromatin template that both permits efficient transcription and prevents inappropriate or spurious transcription. Many of these proteins interact with RNA polymerase II (Pol II) during transcription elongation, when nucleosomes are dynamically modified, disrupted, and reassembled upon Pol II passage [reviewed in (1, 2)]. Results from extensive genetic, biochemical, and structural studies in yeast and mammalian cells support a dynamic interplay in which transcription both regulates and is regulated by chromatin structure. Critical to this interplay are transcription elongation factors, histone chaperones, histone modifiers, and nucleosome remodeling enzymes.

The Polymerase Associated Factor 1 complex or Paf1C is a conserved member of the Pol II elongation complex found on actively transcribed regions of the genome (3–5). *Saccharomyces cerevisiae* Paf1C --- consisting of the five subunits Paf1, Ctr9, Cdc73, Rtf1, and Leo1 --- was first discovered as a Pol II interacting complex engaged in functional and physical interactions with other transcription elongation factors (6–9). Subsequent studies showed that the complex is multi-functional [reviewed in (10)]. *In vitro* transcription reactions with purified factors showed that human Paf1C (hPaf1C) can stimulate transcription elongation on chromatin templates in cooperation with TFIIS (11). More recently, depletion of individual Paf1C subunits, including Paf1 and Rtf1, from human or mouse cell lines has revealed positive effects of the complex on elongation rate and Pol II processivity as well as context-dependent effects on promoter proximal Pol II pausing (12–17). The ability of hRtf1 to stimulate Pol II elongation rate *in vitro* involves an intimate contact between the hRtf1 latch domain and the Pol II catalytic center (5). Mutation or depletion of Paf1C also impairs processes coupled to transcription, including RNA 3’-end formation of noncoding RNAs, polyA site selection at protein-coding genes, mRNA export, and DNA damage repair (18–23).

In addition to directly affecting transcription elongation by Pol II, Paf1C regulates the epigenetic state of transcribed chromatin. In yeast, absence of Paf1, Ctr9, and to a lesser extent, Cdc73, leads to a decrease in H3 lysine 36 trimethylation (H3K36me3) (24). Enriched at the 3’ ends of genes, H3K36me3 promotes histone deacetylation by the Rpd3S complex, culminating in the restoration of a repressed chromatin state following Pol II passage [reviewed in (25)]. Another transcription-associated modification, H2B mono-ubiquitylation on K123 in *S. cerevisiae* (K120 in humans) is strongly dependent on a small conserved domain in Rtf1 (26–28) and is also stimulated by other subunits within the complex, including Ctr9 and Cdc73 (29). The Rtf1 histone modification domain (HMD) interacts directly with the ubiquitin conjugase Rad6 *in vivo* and can stimulate H2BK123ub in a reconstituted *in vitro* reaction (27). H2B123ub initiates a cascade of downstream histone modifications with important roles in transcription, including di- and tri-methylation of H3 K4 and K79 [reviewed in (30)].

Coupling of the many functions of Paf1C to transcription elongation relies on its association with Pol II. Paf1C recruitment to the active Pol II elongation complex is dependent on the phosphorylation state of the Pol II C-terminal domain (CTD of the Rpb1 subunit) as well as the phosphorylation state of the Spt5 C-terminal repeat region (CTR) (31–35). The Spt5-Spt4 complex (DSIF in humans) is a highly conserved component of the active Pol II elongation complex along with Paf1C and Spt6, an essential histone chaperone and elongation stimulatory factor [reviewed in (36–38)]. A combination of genetic, biochemical and structural work has provided a molecular-level understanding of the Paf1C-Spt5 interaction. The central Plus3 domain of Rtf1 binds directly to the phosphorylated CTR of Spt5 (5,39,40). Mutation of the Rtf1 Plus3 domain, the Spt5 CTR, or the yeast CDK9 kinase Bur1, which phosphorylates the Spt5 CTR, greatly reduces Paf1C occupancy on coding regions (31,32,35,39,40). While the importance of the Spt5-Rtf1 interaction for Paf1C recruitment is well supported in yeast, it is unclear if a similar or distinct recruitment mechanism functions in mammalian cells where Rtf1 appears to be transiently associated with the other Paf1C subunits and may not provide a strong attachment point to Pol II (41).

In addition to a recruitment role for the Rtf1 Plus3 domain, we previously demonstrated that the C-terminal domain of yeast Cdc73, which has a Ras-like fold, is important for Paf1C localization to coding regions (42). However, significantly less is known about the interactions between Cdc73 and the Pol II elongation complex. While deletion of the Cdc73 C-terminal domain (hereafter, C-domain) reduced Paf1C occupancy on active genes, it did not affect Pol II occupancy or overall integrity of Paf1C. Biochemical insights into potential interacting partners for Cdc73 came from *in vitro* pulldown assays which detected interactions between Cdc73 and phosphorylated peptides derived from both the Pol II CTD and the Spt5 CTR (33). However, it is unclear if these interactions occur *in vivo* or if Cdc73 contacts other members of the core Pol II elongation machinery. Relevant to this, current structural information on Cdc73 within the context of the active elongation complex is limited (4, 5).

In this study, we used site-specific crosslinking to identify interacting partners for Cdc73 *in vivo* and found that the Cdc73 C-domain directly interacts with two essential components of the Pol II elongation complex, Rpb1 and Spt6. We also demonstrate that Cdc73 can directly interact with Spt6 in the absence of other factors, and we map the binding interface by chemical crosslinking and mass spectrometry. Discovery of the Cdc73-Spt6 interaction led us to investigate a broader role for Spt6 in regulating Paf1C recruitment during transcription elongation using an *spt6* mutant and acute Spt6 depletion. Our results show that Spt6 is critical for establishing and/or maintaining the occupancy of Paf1C and its chromatin-associated functions across the yeast genome.

## MATERIALS AND METHODS

### Yeast strains, growth conditions, and genetic manipulations

*S. cerevisiae* strains are isogenic to FY2, a *GAL2^+^* derivative of S288C (43) and are listed in Supplementary Table S1. Epitope tagging was achieved by one-step integration into the yeast genome and confirmed by PCR (44). Yeast crosses and transformations were conducted as described (45). The strain containing the *rtf1-R251A,Y327A* allele integrated at the *RTF1* locus was created using *delitto perfetto* (46). A region of the *RTF1* gene (+710 to +990) was replaced with pCORE-UH (46) in strain KY3900 by homologous recombination, selecting for hygromycin^R^ Ura*^+^* colonies to generate two independently derived integrants, KY3907 and KY3908. A DNA fragment of *RTF1* containing the R251A and Y327A substitutions was used to replace the pCORE region to generate KY3915 and KY3916. Finally, KY3917 and KY3918, strains that could be used for plasmid shuffling the *spt6* mutations, were made by transformation of KY3915 and KY3916 with pCK25, a *URA3*-marked plasmid containing *FLAG-SPT6* (47), followed by screening for Trp^-^ colonies that had lost pKB1623*. spt6* mutant strains were generated by plasmid shuffling with *TRP1-*marked *CEN/ARS* plasmids expressing 3xV5-tagged WT Spt6 or mutant proteins (48). Transformants were passaged twice on synthetic complete medium (45) lacking tryptophan (SC-W) and containing 0.1% 5-fluoroorotic acid (5-FOA) to select against the *URA3*-marked *SPT6* plasmid. For auxin-induced degradation of Spt6, *SPT6-AID* cells were grown to log phase at 30°C in YPD medium (45) supplemented with 400 µM tryptophan (W). Indole-3-acetic acid (IAA; Sigma-Aldrich, I3750), dissolved in DMSO, was added to the culture at a final concentration of 500 μM before incubation at 30°C for the indicated time prior to crosslinking for chromatin immunoprecipitation (ChIP) or harvesting by centrifugation for western analysis and growth assays.

### Plasmid construction

Plasmid DNA was purified from *E. coli* DH5α cells using a GeneJET Plasmid Miniprep kit (ThermoFisher Scientific K0503). Plasmids were generated by standard restriction digests, site-directed mutagenesis, or Gibson assembly (New England Biolabs E2611S) and verified by DNA sequencing. In plasmid pCK858 (gift from Craig Kaplan), a modification of *RPB1* with a TEV protease cleavage site at amino acid position 1461 is inserted into vector pRS315 (49). Plasmids are listed in Supplementary Table S2 and sequences are available upon request.

### Yeast extract preparation and western analysis

Protein extracts used in western analysis were prepared from log-phase cultures of yeast cells by TCA (50) or post-alkaline extraction (51) or by bead-beating extractions in RIPA or TEV cleavage buffer (50 mM Tris-HCl pH 8.0, 0.5 mM EDTA pH 8.0, 1 mM DTT, 1 mM PMSF) (27). Proteins were resolved on SDS-PAGE gels and transferred to nitrocellulose membranes. Blocking was carried out for 2 hr at room temperature in TBST containing 5% milk.

Antibodies that recognize the following proteins and epitope tags were used: glucose-6-phosphate dehydrogenase (G6PDH; Sigma-Aldrich A-9521, 1:20,000 or 1:50,000), H2B (Active motif 39237, 1:2,000), H2BK123ub (Cell Signaling 5546, 1:1,000), HA tag (Roche 11666606001, 1:1,000 or 1:3,000), His tag (Abcam ab18184, 1:1,000), HSV tag (Sigma-Aldrich H6030, 1:1,000 or 1:2,000), Rpb1-Core (Y80 Santa Cruz, SC-25758, 1:1,000), Rpb1-CTD (8WG16 BioLegend, 664906, 1:500), Rpb3 (BioLegend 665004, 1:1,000), Rtf1 (1:5,000 or 1:2,500 dilution) (9), Spt5 (gift from Grant Hartzog, 1:1,000), Spt6 (gift from Tim Formosa, 1:1,000), and V5 tag (Invitrogen R960-25, 1:1,000).

Primary antibody incubations were carried out overnight at 4°C with agitation. Incubations with anti-mouse (Cytiva NA931) or anti-rabbit (Cytiva NA934) secondary antibodies (1:5,000) were performed at room temperature for 1-2 hr. Visualization of signals was achieved using Pico Plus chemiluminescence substrate (ThermoFisher Scientific 34580) and the ChemiDoc XRS imaging platform (BioRad). Quantifications were performed using ImageJ.

### *In vivo* photocrosslinking

BPA (p-benzoyl-L-phenylalanine) crosslinking experiments were performed as described (27, 52). Briefly, yeast strains BTY42 (Fig. 1 C-E) and KY2838 (Fig. 1F) were co-transformed with pLH157/*LEU2*, the tRNA/tRNA synthetase plasmid for BPA incorporation, and a *TRP1*-marked plasmid harboring *cdc73* with an amber codon substitution at the position of interest to allow for BPA incorporation by nonsense suppression (53). Yeast cells were grown to log phase in SC-L-W medium containing 1 mM BPA (Bachem F-2800). Crosslinking was performed as described (27). For Figures 1C-E, protein extracts were prepared by TCA extraction and analyzed by western blotting. For Figure 1F, 50 mL of cells in log phase were harvested for crosslinking. Following the UV exposure (365 nm), extracts were prepared from cell pellets by bead-beating in 500 µL of TEV cleavage buffer and quantified by Bradford assay. One half of the extract was incubated at 25°C for 1.5 hr with 4 µL of TEV protease (Invitrogen 12575-015). Western blot analysis was performed on 75 µg of extract.

**Figure 1.**
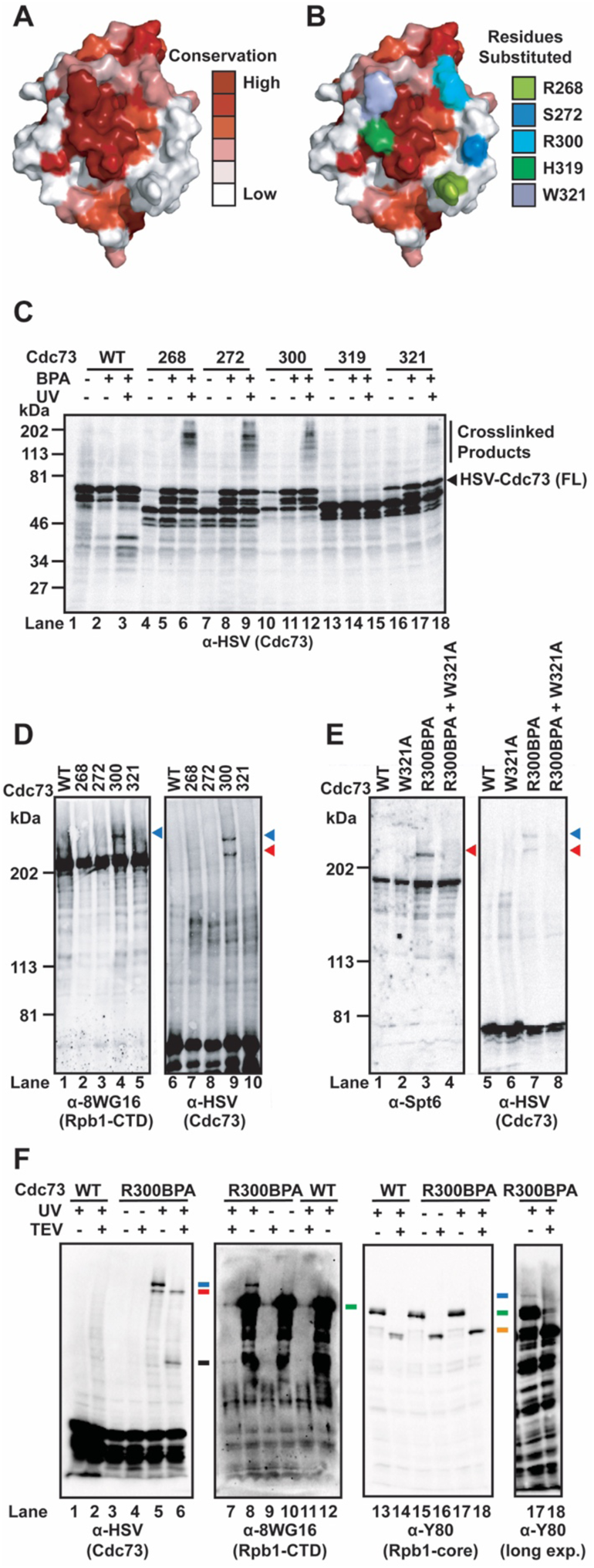
Detection of Cdc73 binding to Spt6 and Rpb1 *in vivo* by BPA crosslinking. (**A**) Cdc73 C-domain crystal structure (3V46) with conservation mapped using the Consurf server. (**B**) As in panel A except showing the locations of residues substituted with BPA. (**C-F**) Western analysis of protein extracts from BPA crosslinking experiments with the indicated antibodies. Cells were grown in the presence of BPA and exposed to UV radiation where indicated. (**C**) Assessment of optimal location for BPA incorporation in N-terminally, triple HSV-tagged Cdc73. Numbers indicate the position of BPA incorporation in Cdc73. On the right, bands corresponding to crosslinked products and full-length HSV-Cdc73 are indicated. Bands below the full-length HSV-Cdc73 band correspond to truncated HSV-Cdc73 proteins that terminate at the amber codon and likely breakdown products. Note, depending on the mutant, we occasionally detect full-length protein in the absence of BPA, indicating endogenous translational readthrough of the amber codon (for example, lane 16). (**D**) Cdc73-Rpb1 interaction (blue arrow) captured by BPA crosslinking. Adjacent panels were derived from the same gel. (**E**) Cdc73-Spt6 interaction (red arrow) captured by BPA crosslinking. Adjacent panels were derived from the same gel. In both panels **D** and **E**, the blue arrow marks the Cdc73-Rpb1 crosslinked product and the red arrow marks the Cdc73-Spt6 crosslinked product. (**F**) Cdc73-Rpb1 interaction analyzed by BPA crosslinking following TEV protease cleavage of the CTD and linker region from the core of Rpb1. Adjacent panels were derived from the same gel. The red line marks the position of the Cdc73-Spt6 crosslinked product, the blue lines mark the full-length Cdc73-Rpb1 crosslinked product prior to TEV cleavage, the green lines mark the position of full-length, uncrosslinked Rpb1, the orange lines mark the position of cleaved, uncrosslinked Rpb1 core domain, and the black line marks the position of the Cdc73-Rpb1 product after TEV cleavage. A longer exposure of lanes 17 and 18 is shown on the right to reveal the Cdc73-Rpb1 crosslinked product (blue line) upon probing with the α-Y80 Rpb1 core domain antibody.

### Recombinant protein expression and purification

Proteins were expressed in *E. coli* CodonPlus RIPL cells using the IPTG induction method. Expression plasmids are listed in Table S2. Cells were grown at 37°C to an OD_600_ of 0.6-0.8 before the addition of 150 µM IPTG and incubation overnight at 25°C. Harvested cell pellets were lysed in Lysis Buffer [Buffer A (25 mM Tris-Cl pH 8.0, 10% glycerol, 1 mM beta-mercaptoethanol (βME)) with 500mM NaCl and 25mM imidazole] by either homogenization or sonication in the presence of a protease inhibitor cocktail containing phenylmethylsulfonyl fluoride, aprotinin, leupeptin, and pepstatin. Lysates were centrifuged at 14,000 rpm for 30 min at 4°C to isolate soluble protein. A combination of nickel affinity chromatography (Qiagen 30210) and cation exchange (GE Healthcare HiTrap-SP; Cdc73) or heparin affinity (GE Healthcare HiTrap-Heparin; Spt6) chromatography was used to purify 10xHis-mClover-Cdc73, Cdc73-10xHis, 10xHis-mClover-Spt6, and 10xHis-mRuby2-Spt6 fusion proteins. For nickel affinity chromatography, Lysis Buffer was used as the loading and wash buffer, and the elution buffer was Buffer A with 100mM NaCl and 250 mM imidazole.

For subsequent ion exchange and heparin affinity chromatography, the NaCl concentration was reduced to 50 mM by dilution with Buffer A prior to column loading, and fractions were taken in 100 mM NaCl steps starting at 100 mM and ending at 1 M. Fractions were collected by passing buffer through the column by hand using a lure lock syringe. Where indicated, the 10xHis tag and accompanying fluorescent protein (mClover or mRuby) were removed by TEV protease digest prior to heparin affinity or cation exchange chromatography. TEV digests were performed overnight, typically at 4°C, in dialysis tubing placed in a minimum of 2 L dialysis buffer (Buffer A with 100mM NaCl). Spt6-239-1451 used in chemical crosslinking and mass spectrometry experiments was further purified by size exclusion chromatography (GE Healthcare Sephacryl S-200) and anion exchange chromatography (GE Healthcare HiTrap-Q). Purified proteins were concentrated and exchanged into Binding Buffer (50 mM HEPES pH 8.0, 100 mM potassium acetate, 5 mM magnesium acetate, 10% glycerol, and 1 mM βME). Buffer exchange was accomplished using Vivaspin concentrators (Sartorius). Protein purity was assessed by SDS-PAGE followed by Coomassie blue staining. A Nanodrop (ThermoFisher Scientific) was used to measure the concentrations of proteins that were not tagged with a fluorophore. Concentrations of fluorescently tagged proteins and any protein used for a quantitative binding assay were determined by SDS-PAGE, using a standard curve (lysozyme or bovine serum albumin) for quantitative comparison.

### Native gel-shift assays

Binding reactions, performed in Binding Buffer (above), contained 150nM 10xHis-mClover-Cdc73 or 10xHis-mClover as a control and a range of concentrations of 10xHis-mRuby2-Spt6 (239-1451) or 10xHis-mRuby2-Spt6 truncation derivatives, as indicated in Figure 2. Reactions were incubated for 10 min at room temperature before loading on the gels. Agarose gels (0.5%) were cast and run using native agarose gel buffer (25 mM Tris-Cl pH 8.0, 19.2 mM glycine) at 120V for 1.5 hrs at 4°C and imaged on an Amersham Imager 600. The gel and buffer were allowed to equilibrate to 4°C for at least 1 hr before gels were loaded. Native gel-shift data were quantified by measuring the intensity of the 10xHis-mClover-Cdc73 band or the 10xHis-mClover control band. Band intensity values were measured from the raw TIFF files produced by the imager using LiCor ImagStudioLite. Fraction shifted was calculated using the following equation: fraction shifted = (shifted band intensity)/(shifted band intensity + unshifted band intensity). Curve fitting was performed with Prism 8 graphing software and the following equation: Y=Bmax*X/(Kd+X)+baseline. Bmax, Kd, and baseline were optimized for curve fitting.

**Figure 2.**
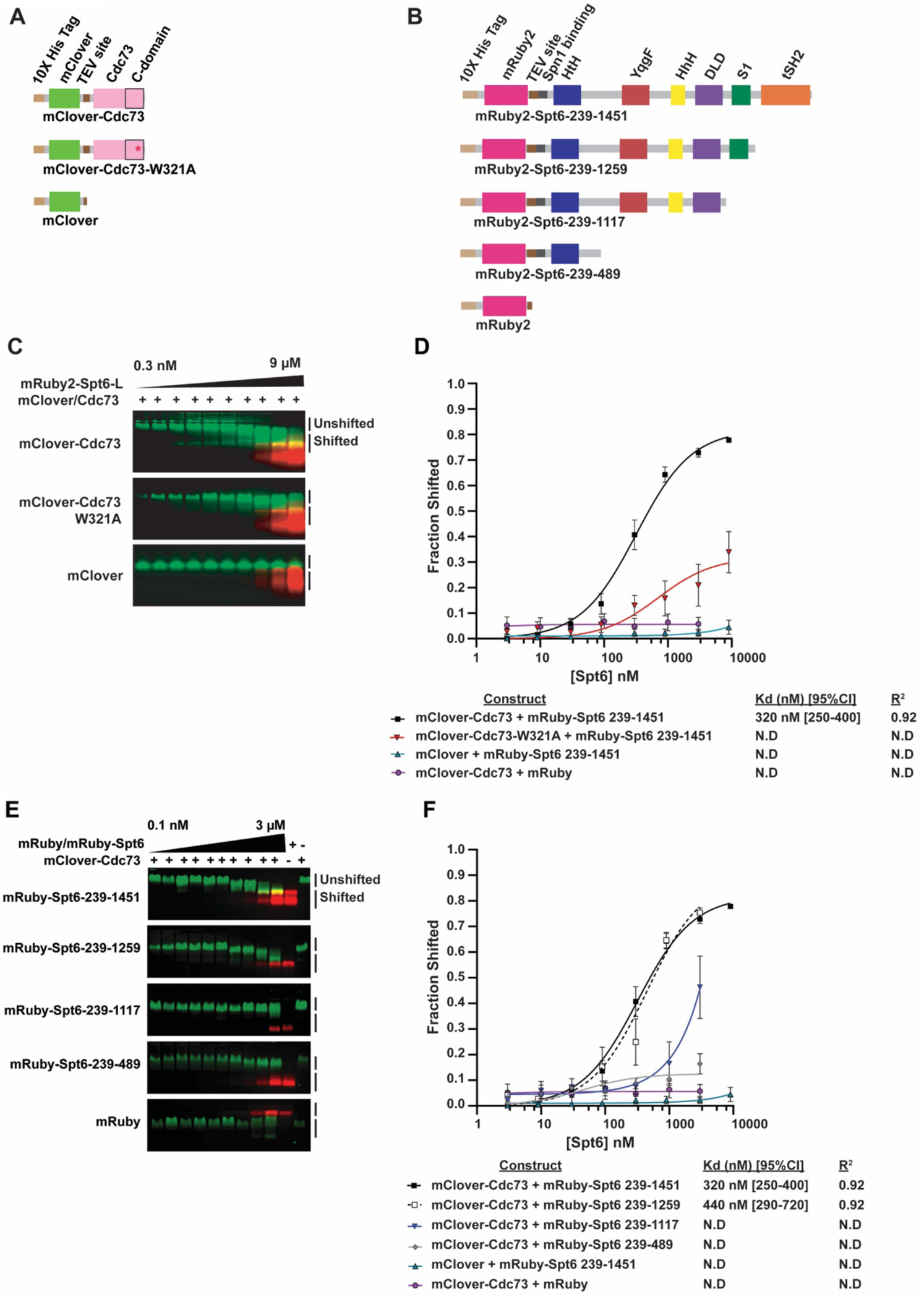
Cdc73 can bind directly to Spt6 *in vitro*. (**A, B**) Diagrams of recombinant, fluorescently tagged proteins used for *in vitro* binding assays. Spt6 protein domain names and locations are as described by Close et al. (65). (**C, E**) Representative native gel-shift assays carried out using recombinantly expressed fusion proteins as diagrammed in panels A and B and shown in Supplementary Figure S1. In panel **C**, mRuby2-Spt6-L (long) denotes the mRuby2-Spt6-239-1451 construct. (**D, F**) Quantified data from the gel-shift experiments shown in panels C and E, respectively, and a minimum of three additional replicates. Data were quantified by calculating the fraction of mClover-Cdc73 (green band) shifted. Binding curves and Kd values were determined in Prism as described in the Methods. N.D. = not determined

### Fluorescence anisotropy

100nM 10xHis-mClover-Cdc73 was mixed with Spt6 (239-1451) in amounts ranging from 750 pM to 7.5 μM in Binding Buffer (see above) in black Nunc 384-Well Polystyrene microplates (ThermoFisher Scientific 262360) at a final reaction volume of 80 μL. Data were collected on a Cytation 5 plate reader (Agilent Technologies) using a green fluorescent polarization filter (excitation and emission wavelengths of 485 nm and 528 nm, respectively) and Gen5 software (BioTek). Curve fitting was performed with Prism 8 graphing software using the following equation: Y=Bmax*X/(Kd+X)+baseline. Bmax, Kd, and baseline were optimized for curve fitting and X was equal to mean change in anisotropy.

### *In vitro* protein crosslinking experiments and mass spectrometry

Crosslinking experiments with DSS (disuccinimidyl suberate, ThermoFisher Scientific 21655) and EDC (1-ethyl-3-(3-dimethylaminopropyl)carbodiimide hydrochloride, ThermoFisher Scientific 22980) were conducted essentially as described with some modifications (54–56). Reactions were performed in a final volume of 20 μL. Proteins were added to either DSS crosslinking buffer/Binding Buffer (50 mM HEPES pH 8.0, 100 mM potassium acetate, 5 mM magnesium acetate, 10% glycerol, and 1 mM βME) or EDC crosslinking buffer (100 mM MES pH 6.0, 100 mM NaCl, 5 mM MgCl_2_, and 10% glycerol) at a final concentration of 10 µM 10xHis-mClover-Cdc73 and 2 µM Spt6 (239-1451). DSS was dissolved in 100% DMSO and added at a final concentration of 625 µM (1 µL of 12.5mM stock), and uncrosslinked controls were treated with 1 µL of 100% DMSO. For EDC crosslinking, EDC and NHS (N-hydroxysuccinimide, ThermoFisher Scientific 24500) were dissolved in water and added at a final concentration of 12.5 mM (EDC, 1 µL of 250 mM) and 250 µM (NHS, 1 µL of 5 mM), respectively, to achieve an EDC:NHS ratio of 50:1. Uncrosslinked control reactions for EDC samples were prepared by addition of 1 µL of water and 1 µL of 5 mM NHS. Reactions were incubated with agitation for 30 min at room temperature and then quenched by addition of 1 µL of 1M Tris-Cl pH 7.5 (DSS) or addition of both 1 µL of 1M Tris-Cl pH 8.0 and 1 µL of 1M BME (EDC), followed by agitation for 15 min at room temperature.

Products from DSS and EDC crosslinking reactions prepared for mass spectrometry analysis were combined with 7 µL of 3X SDS-PAGE loading dye lacking reducing agent, brought to a final concentration of 20 mM DTT by addition of 0.56 µL of 1 M DTT, and incubated at 75°C for 10 min. Samples were allowed to cool to room temperature for 5 min before addition of 1.43 µL iodoacetamide (Sigma A3221-10VL) dissolved in 80% distilled deionized water and 20% acetonitrile (ThermoFisher Scientific 85188) bringing the final concentration to 50 mM (2.5X that of DTT). Samples were incubated at room temperature for 30 min in the dark before running 20 µL of the reaction on an SDS-PAGE gel (4%-12% Bis/Tris; NuPage NP0321). Crosslinked products were verified by western analysis. For mass spectrometry, bands containing crosslinked products were excised from the gel and destained. In-gel digestion of proteins with trypsin and Lys-C and subsequent steps in the LC/MS analysis were performed using a QE HFX Orbitrap mass spectrometer as previously described (55–58). No more than three missing trypsin cleavage sites were allowed. Search results obtained using a 5% false discovery rate were inspected manually to remove potential false positives as previously described (55–58).

### Determining a structural model from crosslinking (XL)/MS data

The xiView web tool was used to generate a two-dimensional network model of the crosslinking data, and structural modeling was performed using the Integrative Modelling Platform (IMP) (59). Prior to modeling with IMP, a prediction of full-length Cdc73 was generated via I-TASSER (60–62). The best prediction of full-length Cdc73 was determined by two metrics: (1) 3D alignment to Cdc73 structures (3V46, 5YDE and 6AF0; (42,63,64)) and (2) the number of valid crosslink lengths by XL-MS that satisfied a 30 Å cutoff. The crosslinking data used when choosing the best prediction were from a separate intramolecular XL-MS experiment on Cdc73-10xHis without Spt6 present (Table S5). This model of *S. cerevisiae* Cdc73 was then used in IMP modeling. Spt6 structural information used with IMP was generated by combining information from 3PSF and 3PSI structures (65) after 3D alignment using the FATCAT server (66–69). This allowed the majority of Spt6 core residues to be included.

The generation of a three-dimensional model was accomplished by IMP using EDC (n=2) and DSS (n=3) datasets, predicted *S. cerevisiae* Cdc73, and Spt6 structural information as input. Crosslinking distances (C_α_ - C_α_) of 20 Å for EDC and 35 Å for DSS were used as constraints for IMP modelling (55). Only inter-protein crosslinks between Cdc73 and Spt6 core were considered in the modeling. During modeling, the position of the Spt6 core was held constant by specifying it as a fixed rigid body, and the Cdc73 structure was allowed to move in order to sample the various Cdc73-Spt6 interaction configurations. The top-scoring models were determined by running the IMP Monte-Carlo (MC) simulation over 100,000 frames with 10 MC steps per frame (1,000,000 steps total). To recover side-chain information, the structural information input into IMP was aligned back to the carbon alpha trace returned by IMP. This step was performed using the alignment plugin in PyMOL. Only Cdc73 data from the 3V46 structure of Cdc73, as defined by the top scoring model, was included in the final model (Figure 3F). Further information on the modeling of the Cdc73-Spt6 interaction, including PDB files, scripts, and crosslinking data, can be accessed on GitHub (https://github.com/mae92/Spt6_Project_XLMS_Structural_Modeling).

**Figure 3.**
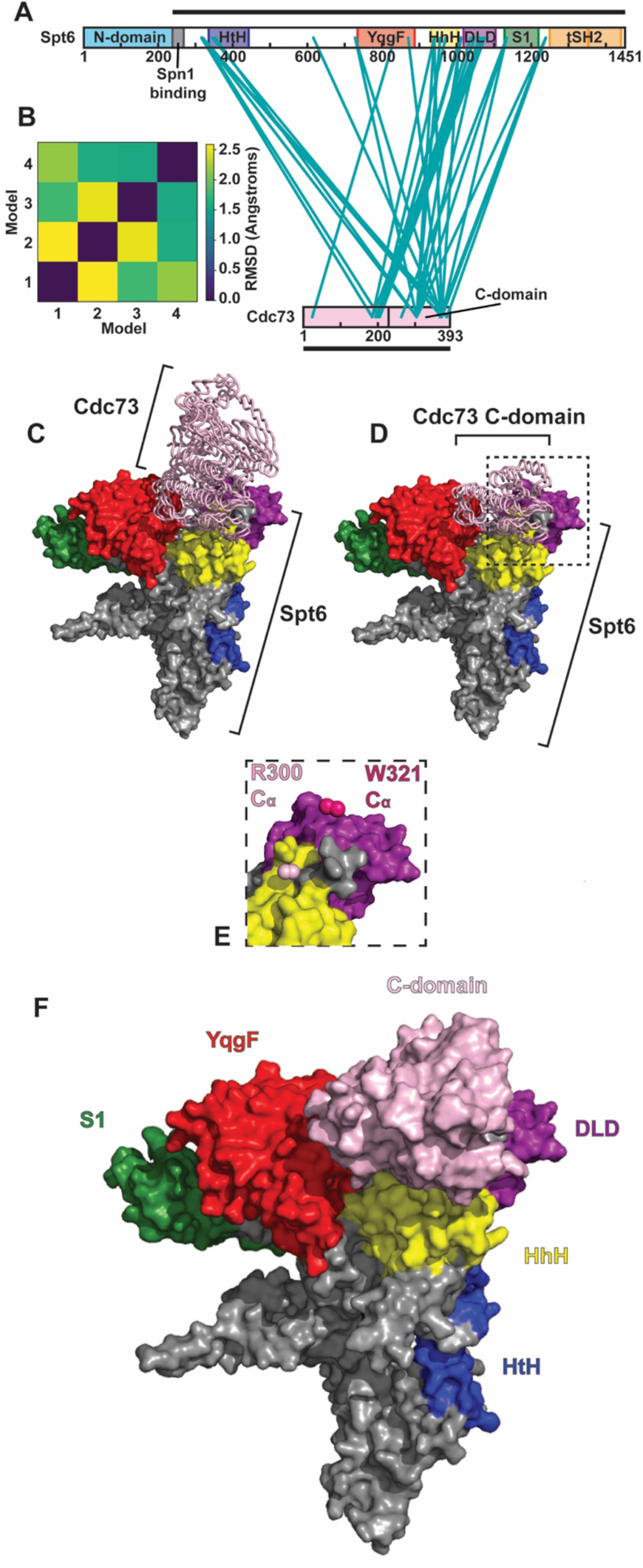
Putative Cdc73-Spt6 interaction interface determined by chemical crosslinking and mass spectrometry. (**A**) Network diagram showing all crosslinks detected between Cdc73 and Spt6 by both EDC and DSS crosslinking methods. Black bars indicate the boundaries of the constructs used for crosslinking. (**B**) Root-mean-square deviation (RMSD) of the top 4 models generated from crosslinking datasets using IMP. (**C, D**) Top 4 structural models of the Cdc73-Spt6 interaction based on crosslinking data. Models show the positions of (**C**) full-length Cdc73 (I-TASSER prediction) and (**D**) the Cdc73 C-domain (both in light pink) on a surface representation of Spt6 (domains colored as in **A**). (**E**) Close-up view showing the positioning of Cdc73 residues R300 (light pink spheres) and W321 (dark pink spheres) relative to the Spt6 surface in the top 4 models. (**F**) Final model of the Cdc73 C-domain (PDB: 3V46) interacting with Spt6 (PDB: 3PSF+3PSI) based on data from the top scoring IMP model.

### Chromatin immunoprecipitation (ChIP)

ChIP was performed essentially as described previously (70, 71). Briefly, 200 or 250 mL of cell cultures were grown to log phase (OD_600_ = 0.8 - 1.0). Cells were crosslinked for 20 min at room temperature by addition of formaldehyde to 1% (v/v). Lysates were prepared in FA lysis buffer (72). Chromatin was sheared in Bioruptor® Pico 15 mL tubes with 300 µL beads (Diagenode C30010017) by 25 cycles of 30 sec on and 30 sec off using a Bioruptor® (Diagenode B01060010). Sonicated chromatin was brought up to a volume of 6 mL in FA lysis buffer and centrifuged at 50,000 rpm for 20 min at 4°C. Chromatin in the supernatant was aliquoted, flash-frozen, and stored at −80°C.

IP reactions were carried out after thawing chromatin on ice and bringing the NaCl concentration to 275 mM. Except for those shown in Fig. S2, 350 µL IP reactions contained chromatin prepared as above and anti-Spt6 (gift from Tim Formosa, 2.5 µL), anti-V5 (Invitrogen R960-25, 1.5 µL), anti-Spt5 (gift from Grant Hartzog, 0.5 µL), anti-HSV (Sigma-Aldrich H6030, 1.25 µL), anti-HA (Santa Cruz sc-7392 AC, 15 µL bead-conjugated antibody), anti-Rtf1 (1 µL; (9)), anti-Rpb3 (BioLegend 665004, 1.25 µL), anti-H2BK123ub (Cell Signaling 5546, 1.25 µL), anti-total H3 (GenScript, 2.5 µL), anti-H3K4me2 (Millipore 07-030, 2.5 µL), anti-H3K4me3 (Active motif 39159, 2.5 µL), or anti-Myc (gift from John Woolford, used as non-specific IgG control, 2.5 µL) primary antibodies. For the ChIP experiment in Fig. S2, 225 µg of *S. cerevisiae* chromatin was mixed with 25 µg of *Schizosaccharomyces pombe* chromatin (KP07 or KP08) in a 350 µL reaction as measured with a Pierce BCA protein assay kit (Thermo Fisher Scientific 23227), and IP reactions were conducted with anti-V5 (Invitrogen R960-25, 1 µL), anti-Rpb1-CTD (8WG16; BioLegend 664906, 1 µL) or anti-HA (Santa Cruz sc-7392 AC, 15 µL bead-conjugated antibody) primary antibodies. For the anti-HSV IP reactions (Sigma-Aldrich H6030, 1.75 µL) in Fig. S2, 450 µg of *S. cerevisiae* chromatin and 50 µg of *S. pombe* chromatin were used. While *S. pombe* chromatin was included for these ChIP assays, it was not used for normalization of the *S. cerevisiae* data as it increased the variance in the measurements. IP reactions were incubated overnight at 4°C on an end-over-end roller. Protein A or G beads (Cytiva 17-5280-01 or 17-0618-01) were washed with FA lysis buffer with 275 mM NaCl and added to the chromatin-antibody mixture and placed on the roller for 1 hr at room temperature. Beads were washed and DNA recovered as previously described (71). DNA was purified using the QIAquick PCR Purification Kit (Qiagen, 28106).

### Quantitative polymerase chain reaction (qPCR)

All qPCR experiments were performed in biological triplicate and technical duplicate using qPCRBIO SyGreen Blue 2x reaction mix (Genesee Scientific 17-505B) and analyzed on a QuantStudio3™ Real-Time PCR System (Thermo Fisher). Efficiencies were determined for primer sets by measuring C_t_ values across a series of six ten-fold dilutions of *S. cerevisiae* genomic DNA. ChIP-qPCR data were analyzed as described (73) and normalized as indicated in the Figure Legends. Primer sequences and efficiencies are listed in Table S3.

### Next-generation sequencing library preparation

ChIP-sequencing (ChIP-seq) libraries were prepared using a NEBNext Ultra II kit and indices (E7645, E7335, E7500, E7710, E7730). Library build reactions contained immunoprecipitated DNA at 0.9 ng/µL and 0.1 ng/µL of spike-in DNA from *K. lactis*. Addition of spike-in DNA by this method controls for library build and sequencing effects but does not allow detection of global changes in protein occupancy. To detect global changes in protein occupancy, we normalized IP to input data via the NCIS method (74). Libraries were quantified by Qubit and fragment length distribution was assessed by agarose gel electrophoresis. DNA sequencing was conducted by the University of Pittsburgh Health Sciences Sequencing Core at UPMC Children’s Hospital of Pittsburgh on an Illumina NextSeq 500.

### ChIP-seq data analysis

ChIP-seq reads were aligned to the *K. lactis* genome (Ensembl ASM251v1) and then unaligned reads were aligned to the *S. cerevisiae* genome (Ensembl R64-1-1), using HISAT2 (75) (options --no-unal –no-mixed –no-discordant –nospliced-alignment –no-softclip -k 2) before SAM to BAM conversion, PCR duplicate removal and subsetting with the SAMtools suite (76). The number of reads subsampled from each BAM file was based on the file with the lowest number of sequenced reads and rounded to the nearest hundred thousand. For the *SPT6* vs *spt6-50* experiments, all data sets were subsampled to 9,000,000 reads. Subsamples for the *SPT6-AID* IAA vs DMSO experiments were as follows: Spt6, HSV-Paf1, Spt5, and Rpb3 (5,000,000); H2BK123ub (6,000,000); H3 (1,800,000); H3K4me2 (1,000,000); and H3K4me3 (1,800,000). Subsampled BAMs were then sorted via SAMtools before normalization relative to input and BigWig conversion with a custom NCIS normalization script. BAM to BigWig conversion was conducted while simultaneously applying the scaling factor generated by the NCIS method (74). This was done by multiplying each basepair of the genome by the inverse of the scaling factor (r, described below) using deepTools3 bamCoverage (options: --scalingFactor = (1/r) –extendReads --binSize = 1).

The NCIS normalization value (r) was calculated for each dataset independently in a data adaptive manner. Briefly, each dataset was broken into 100 bp bins and the ratio of IP/Input was calculated for each bin. The correct number of bins (n) to use in calculating r was determined by titrating the per bin read count cutoff (t; based on IP sample). In this process t is increased by 1 in an iterative manner until r is greater than or equal to the r calculation from the last iteration and the pool of normalization bins accounted for 2% of the genome. At this point, the sum of n bins with t or less reads measured for the IP sample is taken for both IP and input datasets and the ratio of IP/Input is calculated from these sums resulting in our normalization factor r.

Biological replicate BigWig files were averaged using the bigwigCompare command (options: --pseudocount 0.1 --operation mean --binSize 1) and log2 fold change BigWigs were generated from these using the bigwigCompare command (options: --operation log2 --binSize 1). Heatmaps and aggregation plots were plotted over genic regions using a bin size of 25 bp and averaging data within each bin using deepTools3 and a combination of the computeMatrix and either the plotHeatmap or plotProfile commands (77, 78). Differential occupancy analysis was conducted in RStudio. ChIP-seq reads were counted from BAM files for all protein-coding genes in the *S. cerevisiae* genome using the featureCounts function in the Rsubread package and used to generate count tables for DESeq2. Count tables were filtered prior to statistical analysis with DESeq2 to only include genes where the sum of the read counts across samples to be analyzed was greater than or equal to 100 reads. The NCIS normalization factor r was applied to all datasets during DESeq2 normalization. The DESeq2 and psych R packages were used to generate statistics on differential ChIP-seq occupancy presented in MA and correlation plots, respectively. Bash and R code is stored on GitHub (https://github.com/mae92/Spt6_Project_ChIPseq_Analysis).

### Statistical analysis and reproducibility

BPA crosslinking experiments were performed at least twice using independent transformants. For ChIP-qPCR, three biological replicates were assayed in technical duplicate. A minimum of three biological replicates were analyzed for the AID western blot and viability assay experiments. Each biological replicate is a pure yeast culture derived from a single colony. For ChIP analysis of *spt6 rtf1* double mutants in Fig. S2, two independent biological replicates were created from KY3917 and one from KY3918 through plasmid shuffling. Protein binding assays were performed at least four times. DSS and EDC crosslinking and mass spectrometry were performed in triplicate and duplicate, respectively, generating a total of five crosslinking datasets for model generation. Bar graphs plot mean and either standard error of the mean or standard deviation (see Figure Legends) and include all individual data points. For ChIP-qPCR data, p-values were generated using an unpaired, two-sided Student’s t-test, assuming equal variance carried out between the mutant strain and the wild-type strain. Pearson’s correlation analysis was used to assess replicate reproducibility and correlation between ChIP-seq datasets. Spearman’s correlation analysis was used to compare log2 fold change datasets. Differential gene occupancy was determined using a negative binomial generalized linear model, which was implemented in R via the DESeq2 package.

## RESULTS

### Cdc73 directly interacts with Rpb1 and Spt6 *in vivo*

To identify interaction partners for Cdc73 *in vivo*, we employed a site-specific protein crosslinking strategy. Guided by our structural analysis of Cdc73 (42), we replaced surface-exposed amino acids on the Cdc73 C-domain (Fig. 1A-B) with the photoreactive phenylalanine analog, BPA, through amber codon suppression (53). Based on the levels of full-length Cdc73 protein produced in cells grown in BPA-containing medium, BPA was incorporated at positions 268, 272, 300 and 321 while incorporation at position 319 was unsuccessful (Fig. 1C). BPA-dependent, crosslinked products were produced upon exposure of cells to UV light at 365 nm, and these were detected by western blotting using an antibody against the HSV tag on Cdc73 (Fig. 1C, lanes 6, 9, 12 and 18). Further resolution of the crosslinked species revealed two slowly migrating, HSV-reactive bands in the R300BPA sample. Western analysis identified the top band as a crosslinked product between HSV-Cdc73 and the largest Pol II subunit, Rpb1 (Fig. 1D, lanes 4 and 9; blue arrow), and the bottom band as HSV-Cdc73 crosslinked to Spt6 (Fig. 1E, lanes 3 and 7; red arrow).

To investigate the functional significance of the Cdc73-Rpb1 and Cdc73-Spt6 interactions, we performed BPA crosslinking experiments with an HSV-Cdc73 R300BPA derivative in which the highly conserved, surface exposed tryptophan at position 321 was changed to alanine. We previously showed that the Cdc73 W321A substitution causes phenotypes associated with defects in transcription elongation (42). Crosslinking of both Rpb1 and Spt6 to HSV-Cdc73 R300BPA was greatly diminished by the W321A substitution suggesting that these interactions are dependent on a transcriptionally important residue within Cdc73 (Fig. 1E).

In previous studies, recombinant full-length Cdc73 and a C-domain containing fragment of Cdc73 exhibited binding to diphosphorylated Pol II CTD peptides *in vitro* (33). Our crosslinking data demonstrate that Cdc73 interacts directly with Rpb1 in cells. To further characterize this interaction and begin to map the site of interaction on Rpb1, we performed BPA crosslinking experiments on yeast cells expressing HSV-Cdc73 R300BPA and a derivative of Rpb1 containing a TEV protease cleavage site inserted between amino acids 1461 and 1462. The TEV cleavage site separates the Rpb1 core domain from the Rpb1 linker region and CTD (79). Following TEV treatment of extracts prepared from UV-exposed cells, the Cdc73-Rpb1 crosslinked product (Fig. 1F, lane 5, blue line) shifted to a position on the gel (Fig. 1F, lane 6, black line) that was inconsistent with crosslinking to the Rpb1 core domain (Fig. 1F, lanes 14, 16, and 18, orange line denotes the position of uncrosslinked Rpb1 core domain). Moreover, the TEV-cleaved Cdc73-Rpb1 product (Fig. 1F, lane 6, black line) was detected with 8WG16 antibody, which recognizes the Pol II CTD (Fig. 1F, lane 7, black line). Detection of the HSV- and 8WG16-reactive band (Fig. 1F, lanes 6 and 7, black line) was dependent on the presence of BPA and on the UV treatment. Together, these results provide the first demonstration that Paf1C subunit Cdc73 directly contacts the Pol II CTD/linker region and Spt6 *in vivo*.

### Cdc73 and Spt6 can interact directly *in vitro*

Because an interaction between Cdc73 and Spt6 had not been previously characterized, we focused on this interaction and a possible role for Spt6 in Paf1C recruitment to transcribed genes. To test if the Cdc73-Spt6 interaction can occur in the absence of other factors, we conducted *in vitro* binding assays with purified, recombinant Spt6 (239-1451) and full-length Cdc73 fused to 10xHis-mRuby2 and 10xHis-mClover tags, respectively (Fig. 2A-B and Fig. S1A). The Spt6 (239-1451) derivative lacks an acidic, unstructured N-terminal region and has been used previously for structural studies (65). Native gel-shift assays, followed by detection and quantification of fluorescence signals, showed that the tagged Spt6 (239-1451) and Cdc73 proteins can interact *in vitro* with a K_d_ of approximately 320 nM (Fig. 2C-D).

Fluorescence anisotropy data collected using 10xHis-mClover-Cdc73 and unlabeled Spt6 (239-1451) support the gel-shift results but reported a lower binding affinity (K_d_ = 770 nM, Fig. S1B). Consistent with our *in vivo* BPA crosslinking results, the W321A substitution in Cdc73 weakened the interaction between recombinant Cdc73 and Spt6 (Fig. 2C-D). These data show that Cdc73 and Spt6 are able to interact in the absence of Pol II and other components of the transcription elongation complex.

To begin to map the site of Cdc73 interaction within Spt6, we purified C-terminally truncated derivatives of 10xHis-mRuby2-Spt6 that lack defined structural domains of the protein (Fig. 2B and Fig. S1A) (65). As measured by native gel-shift assays, removal of the Spt6 tandem SH2 (tSH2) domains had no effect on the Cdc73-Spt6 interaction, but removal of the tSH2 domains and the S1 domain greatly reduced the interaction with 10xHis-mClover-tagged Cdc73 (Fig. 2E-F). An Spt6 truncation mutant lacking all structural domains C-terminal to the helix-turn-helix domain (HtH) was severely defective for Cdc73 binding (Fig. 2E-F). These results indicate that the Spt6 core, which extends from the HtH domain through the S1 domain (65), is important for the interaction between Cdc73 and Spt6 *in vitro*.

To elucidate the interface between recombinant 10xHis-mClover-Cdc73 and untagged Spt6 (239-1451), we performed chemical crosslinking followed by mass spectrometry. For this analysis, we used two different crosslinking agents: DSS, which crosslinks lysine residues, and EDC, which crosslinks lysines to either glutamic acid or aspartic acid residues (55). Proteins exposed to crosslinking agents were resolved by SDS-PAGE, and bands representing the 1:1 complex, as judged by mass and western analysis, were excised and analyzed by mass spectrometry. Interprotein crosslinks were observed between Cdc73 and several conserved domains within the Spt6 core (Fig. 3A, Table S4). The most heavily crosslinked domains of Spt6 were the death-like domain (DLD) and helix-hairpin-helix (HhH) domain (Fig. 3A and Table S4). With respect to Cdc73, most of the crosslinks mapped to the C-domain (between residues 263 and 371) and the region N-terminal to the C-domain (between residues 185 and 205). These observations are consistent with our *in vivo* BPA crosslinking results (Fig. 1) and with a recent affinity purification-mass spectrometry study, which revealed significantly diminished Paf1C association with Pol II in yeast cells harboring the *spt6-1004* mutation, which deletes the HhH domain and reduces Spt6 levels (80).

We used interprotein crosslinks from replicate experiments, Spt6 structural information, and a predicted model of full-length Cdc73 (see Methods and Tables S4 and S5) as input for the Integrative Modeling Platform (IMP) (59). The top 4 structural models produced by IMP aligned with an RMSD <2.5 Å (Fig. 3B). The top-scoring model showed Cdc73 interacting with the Spt6 core in a depressed region between the DLD, HhH, and YqgF domains (Fig. 3C-F and Table S4). Consistent with results from our BPA crosslinking experiments, residues W321 and R300 face in toward the Spt6 core in this model (Fig. 3D-E).

Guided by our structural model (Fig. 3F), we generated alanine scanning substitutions in Spt6 that map within the predicted Cdc73-Spt6 interface and tested their effects on Paf1C occupancy at a highly transcribed gene, *PMA1*, *in vivo* (Fig. S2). The substitutions were designed to change three to five consecutive residues to alanines within or near the YqgF (NRK_771-773_ to AAA), HhH (NKATD_936-940_ to AAAAA), and DLD (KRQK_1004-1007_ to AAAA; ELLRE_1058-1062_ to ALLAA) domains of Spt6 (Fig. S2A-B). *CEN/ARS* plasmids harboring the *spt6* mutations were introduced into an *spt6Δ* strain by plasmid shuffling. Western analysis demonstrated that the V5-tagged Spt6 mutant proteins were present at a similar level as wild-type Spt6 and did not affect the levels of Rpb1 or three different subunits of the Paf1 complex, Paf1, Cdc73 and Rtf1 (Fig. S2C). The ChIP-qPCR analysis showed that none of the individual alanine scanning substitutions significantly impacted Spt6 or Paf1C occupancy at *PMA1*, although slight effects on Rpb1 occupancy were observed (Fig. S2D). To test if the putative Cdc73-Spt6 interface functions redundantly with the Rtf1-Spt5 interaction in recruiting Paf1C to chromatin, we introduced the *spt6* mutations into an *rtf1* mutant background, *rtf1-R251A, Y327A*, in which residues at the interface between the Rtf1 Plus3 domain and the Spt5 CTR are altered (40). While the *rtf1-R251A, Y327A* mutation decreased both Paf1 and Cdc73 occupancy at *PMA1* as expected (40), addition of the *spt6* mutations did not exacerbate this effect (Fig. S2D). These observations suggest that the proposed Cdc73-Spt6 interface is either not playing a major role in Paf1C recruitment at *PMA1* or the individual *spt6* alanine-scanning mutations are insufficient to disrupt the Cdc73-Spt6 interaction.

### An *spt6* mutation that disrupts the Pol II-Spt6 interaction reduces Paf1C occupancy on gene bodies

Reasoning that the targeted substitutions at the predicted Cdc73-Spt6 interface might be insufficient to disrupt the Spt6-Paf1C interaction, we investigated a broader role for Spt6 in controlling Paf1C occupancy at transcribed genes. To this end, we performed ChIP-seq experiments on *SPT6* and *spt6-50* strains in biological duplicate, measuring occupancy of Rpb3, Spt5, Spt6, and HSV-tagged Paf1 (see Fig. S3 for replicate comparisons). The *spt6-50* mutation is a nonsense mutation that truncates the Spt6 tSH2 domain (65,79,81–84) (Fig. 4A), which interacts directly with the Rpb1 linker (79) and is important for coupling Spt6 to Pol II on gene bodies (3, 79). Therefore, if Spt6 is important for recruiting or retaining Paf1C on transcribed genes, we expected a decrease in Paf1 occupancy at genes where Spt6 occupancy is reduced in the mutant. ChIP-seq data obtained for 3087 non-overlapping protein-coding genes were normalized by the NCIS (<u>N</u>ormalization of <u>C</u>hIP<u>s</u>eq) method (74). The NCIS method enables calculation of a normalization factor for each dataset independently in a data adaptive manner. Western analysis showed that total protein levels for Spt6, HSV-Paf1, Rtf1, and Spt5 were similar in the *SPT6* and *spt6-50* strains; however, Rpb3 levels were moderately reduced in the presence of HSV-tagged Paf1 (Fig. 4B).

**Figure 4.**
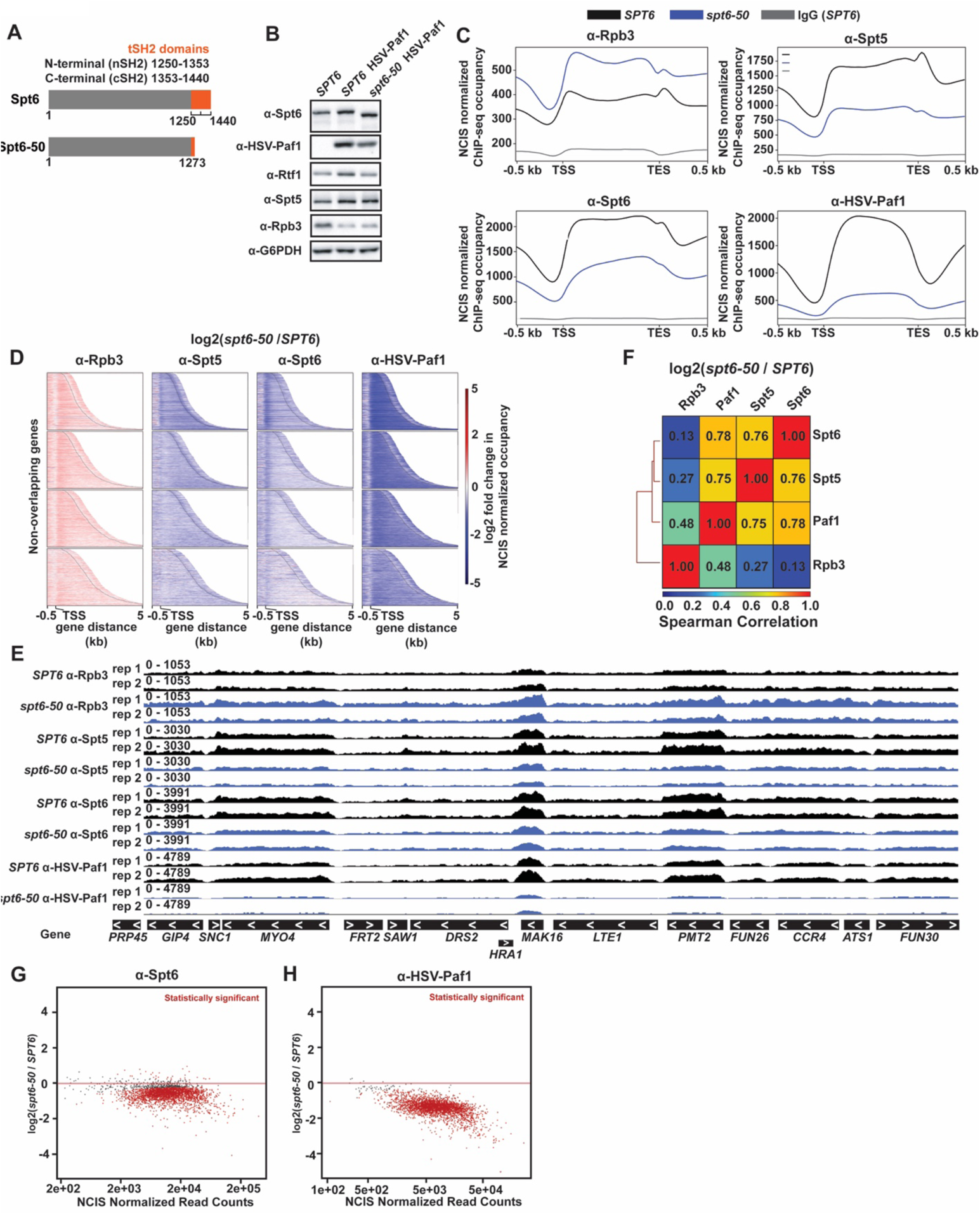
Disruption of the Pol II-Spt6 interaction via the *spt6-50* mutation leads to a reduction in Paf1 occupancy. (**A**) Diagram of the proteins encoded by the *SPT6* and *spt6-50* genes. The *spt6-50* mutation replaces codon 1274 with a stop codon (Fred Winston, personal communication). (**B**) Western analysis showing levels of proteins analyzed by ChIP. G6PDH serves as the loading control. (**C**) Aggregation plots of Rpb3, Spt5, Spt6, and HSV-Paf1 NCIS-normalized ChIP-seq occupancy. IgG was used as a background control for ChIP-seq with chromatin prepared from the *SPT6* strain. (**D**) Heatmaps of differential Rpb3, Spt5, Spt6, and HSV-Paf1 occupancy (*spt6-50* vs *SPT6*) with genes divided into quartiles based on Rpb3 occupancy and sorted by length. Heatmaps show the region between the transcription start site (TSS) and the cleavage and poly-adenylation site (labeled with a black line) +/- 500 bp and use a bin size of 25 bp. (**E**) Genome browser tracks of a region of chromosome 1 (chrI:84,656-118,640) showing NCIS-normalized ChIP-seq occupancy for individual biological replicates. (**F**) Heatmap of Spearman correlation between datasets plotted in D. (**G, H**) MA plots showing differential occupancy for Spt6 and HSV-Paf1 determined by DESeq2 (*spt6-50* vs *SPT6*). Genes with statistically significant changes in occupancy (FDR-adjusted p value ≤ 0.05) are colored red. The results presented in panels C, D, F, G, and H were generated by averaging NCIS-normalized ChIP-seq data from two biological replicates for 3087 non-overlapping protein-coding genes (92).

The NCIS-normalized ChIP-seq data revealed a genome-wide reduction in Spt6 occupancy in the *spt6-50* mutant as expected (3), despite a slight increase in Rpb3 occupancy (Fig. 4C, D, G; Fig. S3B and S3D; Fig. S4C). The reduction in Spt6 occupancy was accompanied by a reduction in occupancy of both HSV-Paf1 and Spt5 with stronger effects observed at genes with the highest levels of Rpb3 occupancy in the wild type (Fig. 4C, D, H; Fig. S3C and S3E; Fig S4D). Browser tracks revealed the changes in protein occupancy at individual genes (Fig. 4E). Consistent with the ChIP-seq data, ChIP-qPCR results, normalized to Rpb3 occupancy, revealed a greater effect of the *spt6-50* mutation on Spt6 and HSV-Paf1 occupancy at highly expressed genes (Fig. S4A) than at lowly expressed genes (Fig. S4B). Changes in Rtf1 occupancy mirrored those of Paf1 (Fig. S4A-B), suggesting that the occupancy of Paf1C, and not Paf1 alone, is reduced in the *spt6-50* mutant. At the genes analyzed by ChIP-qPCR, the impact of the *spt6-50* mutation on Spt5 occupancy was less apparent (Fig. S4A-B).

Correlation analysis of the ChIP-seq data revealed that the effect of the *spt6-50* mutation on HSV-Paf1 occupancy was more highly correlated with the effects of the mutation on Spt6 and Spt5 than Rpb3 (Fig. 4F). These data suggest that disruption of the Pol II-Spt6 interaction via the *spt6-50* mutation leads to a reduction in Paf1 occupancy on gene bodies, supporting a role for Spt6 in recruiting or retaining Paf1C on transcribing Pol II. However, because our ChIP-seq analysis also revealed a reduction in Spt5 occupancy in the *spt6-50* mutant, these results do not allow us to disentangle the impact of Spt6 on Paf1C occupancy from that of Spt5, which is known to play a key role in Paf1C recruitment through its interaction with Rtf1 (32,35,39,40).

### Acute depletion of Spt6 leads to genome-wide loss of Paf1 occupancy and changes in histone modification patterns

To test more directly if Spt6 impacts Paf1C occupancy, we acutely depleted Spt6 through an auxin-inducible degron tag (85) appended to the C-terminus of Spt6, followed by ChIP-seq analysis of HSV-tagged Paf1 (Fig. 5A). This approach allowed us to assess the effects of rapidly depleting Spt6, while minimizing cellular adaption to its loss and indirect effects. A viability assay confirmed that treatment of the *SPT6-AID* strain with auxin for one hour did not result in significant cell death relative to a DMSO-treated control sample (Fig. S5A). Western analysis demonstrated a reduction in Spt6 levels over a one-hour time course of auxin treatment with no adverse effect on HSV-Paf1 levels (Fig. 5B and Fig. S5B-C).

**Figure 5.**
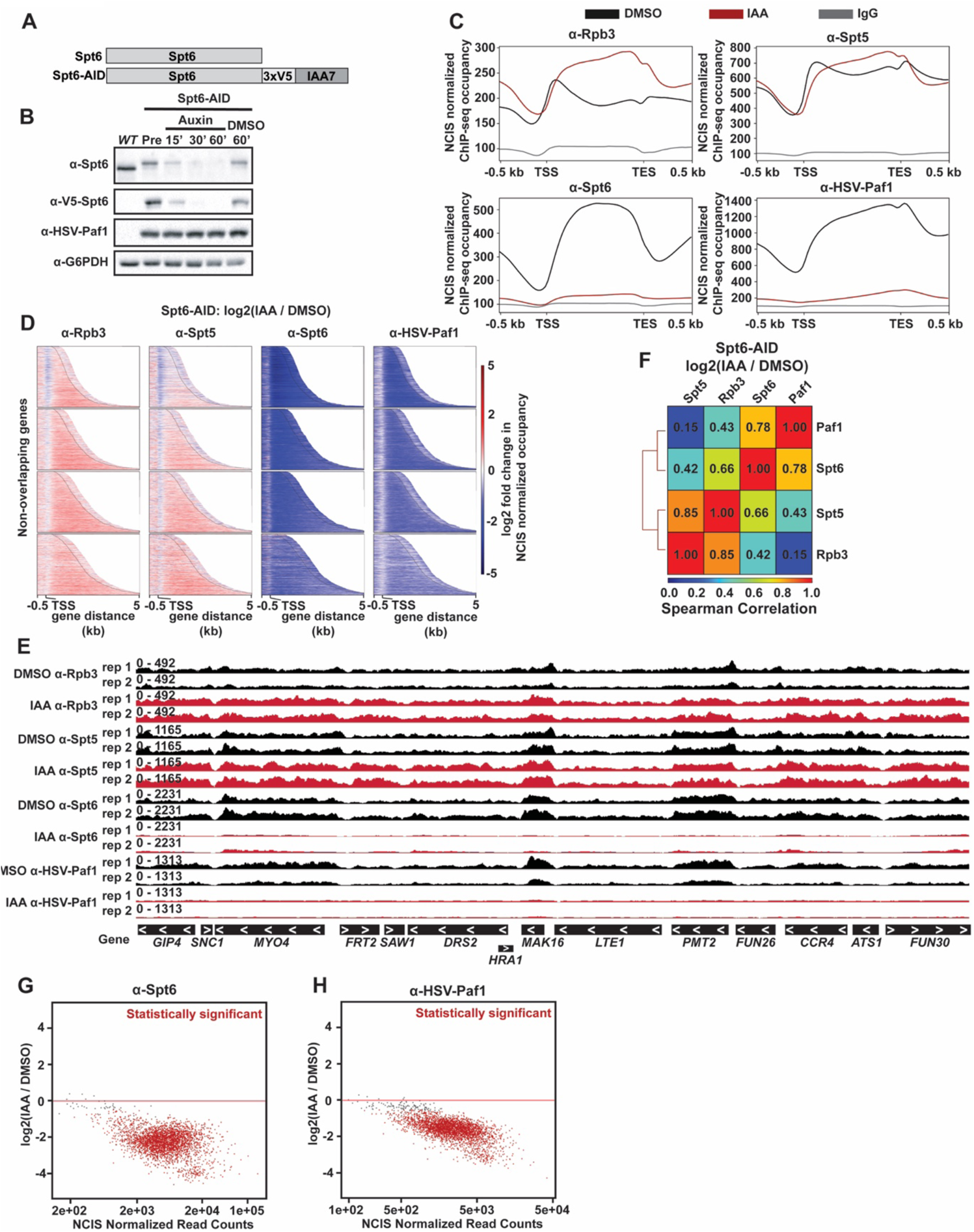
Acute depletion of Spt6 leads to nearly complete loss of Paf1 occupancy genome-wide. (**A**) Diagram comparing the proteins encoded by the *SPT6* and *SPT6-AID* genes. (**B**) Western blot of yeast cells exposed to IAA for 0 (Pre), 15, 30, or 60 min or DMSO for 60 min. An *SPT6* strain (WT), lacking tags on *SPT6* and *PAF1*, was used as a control to measure abundance of the Spt6-AID protein prior to depletion. Levels of G6PDH serve as a loading control. (**C**) Aggregation plots of Rpb3, Spt5, Spt6, and HSV-Paf1 NCIS-normalized ChIP-seq occupancy. IgG was used as a background control for ChIP-seq. (**D**) Heatmaps of differential Rpb3, Spt5, Spt6, and HSV-Paf1 occupancy (IAA vs DMSO) with genes divided into quartiles based on Rpb3 occupancy and sorted by length. Heatmaps show the region between the transcription start site (TSS) and the cleavage and poly-adenylation site (labeled with a black line) +/- 500 bp and use a bin size of 25 bp. (**E**) Genome browser tracks of a region of chromosome 1 (chrI:84,656-118,640) showing NCIS-normalized ChIP-seq occupancy for individual biological replicates. (**F**) Heatmap of Spearman correlation between datasets plotted in D. (**G, H**) MA plots showing DESeq2 differential occupancy results for Spt6 and HSV-Paf1 NCIS-normalized ChIP-seq occupancy (IAA vs DMSO). Genes with statistically significant changes in occupancy (FDR-adjusted p value ≤ 0.05) are colored red. The results presented in panels C, D, F, G, and H were generated by averaging NCIS-normalized ChIP-seq data from two biological replicates for 3087 non-overlapping protein-coding genes (92).

As measured by NCIS-normalized ChIP-seq analysis of two biological replicates, *SPT6-AID* cells treated with auxin for one hour showed a dramatic and statistically significant decrease in both Spt6 and HSV-Paf1 occupancy on chromatin compared to *SPT6-AID* cells treated with DMSO (Fig. 5C-H; see Fig. S6 for correlations between samples and between replicates). In contrast to the effects on HSV-Paf1, rapid depletion of Spt6 caused a slight increase in both Rpb3 and Spt5 occupancy over gene bodies (Fig. 5C-E, Fig. S5F and S5G), potentially due to increased internal transcription initiation events and/or reduced elongation rate in the absence of Spt6 (86–88). Correlation analysis of changes in occupancy profiles upon Spt6 depletion separated Spt5 and Rpb3 into one group and Paf1 and Spt6 into another (Fig. 5F). In support of the ChIP-seq data, ChIP-qPCR experiments measuring the occupancies of Spt6 and Paf1 relative to Rpb3 at highly and lowly expressed genes revealed a significant decrease in Spt6 occupancy at both gene classes and a significant decrease in Paf1 occupancy at a highly expressed gene (Fig. S5D-E). These data implicate Spt6 as a critical factor in controlling proper genome-wide localization of Paf1C in yeast.

Given the importance of Paf1C in promoting transcription-coupled histone modifications, we assessed the effects of rapidly depleting Spt6 on the genic patterns of H2BK123ub, H3K4me2 and H3K4me3 (see Fig. S7 for correlations between samples and between replicates). In agreement with previous studies on *spt6* mutant strains (89–91), a global reduction in H3 occupancy was observed when *SPT6-AID* cells were treated with auxin for one hour (Fig. 6A-D). In addition to the general reduction in H3 occupancy, we observed a significant loss of H2BK123ub across gene bodies and a redistribution of H3K4me2 and H3K4me3 toward the 3’ ends of genes (Fig. 6A-D). The reduction in H2BK123ub correlates with the loss of Paf1 occupancy observed upon Spt6 depletion (Fig. 5; Fig. S8). The changes in H3K4me2 and H3K4me3 patterns upon Spt6 depletion correlate with one another (Fig. S8) and strongly agree with the results of Jeronimo *et al*. (90), who observed a 3’-directed redistribution of these marks in *spt6-1004* mutant cells or upon anchor-away depletion of Spt6. Redistribution of H3K4me3 was also observed upon acute depletion of Spn1, a histone chaperone that interacts with Spt6 (92). By demonstrating a requirement for Spt6 in the recruitment and/or retention of Paf1C on gene bodies, our results provide an additional mechanism through which Spt6 determines histone modification patterns genome-wide.

**Figure 6.**
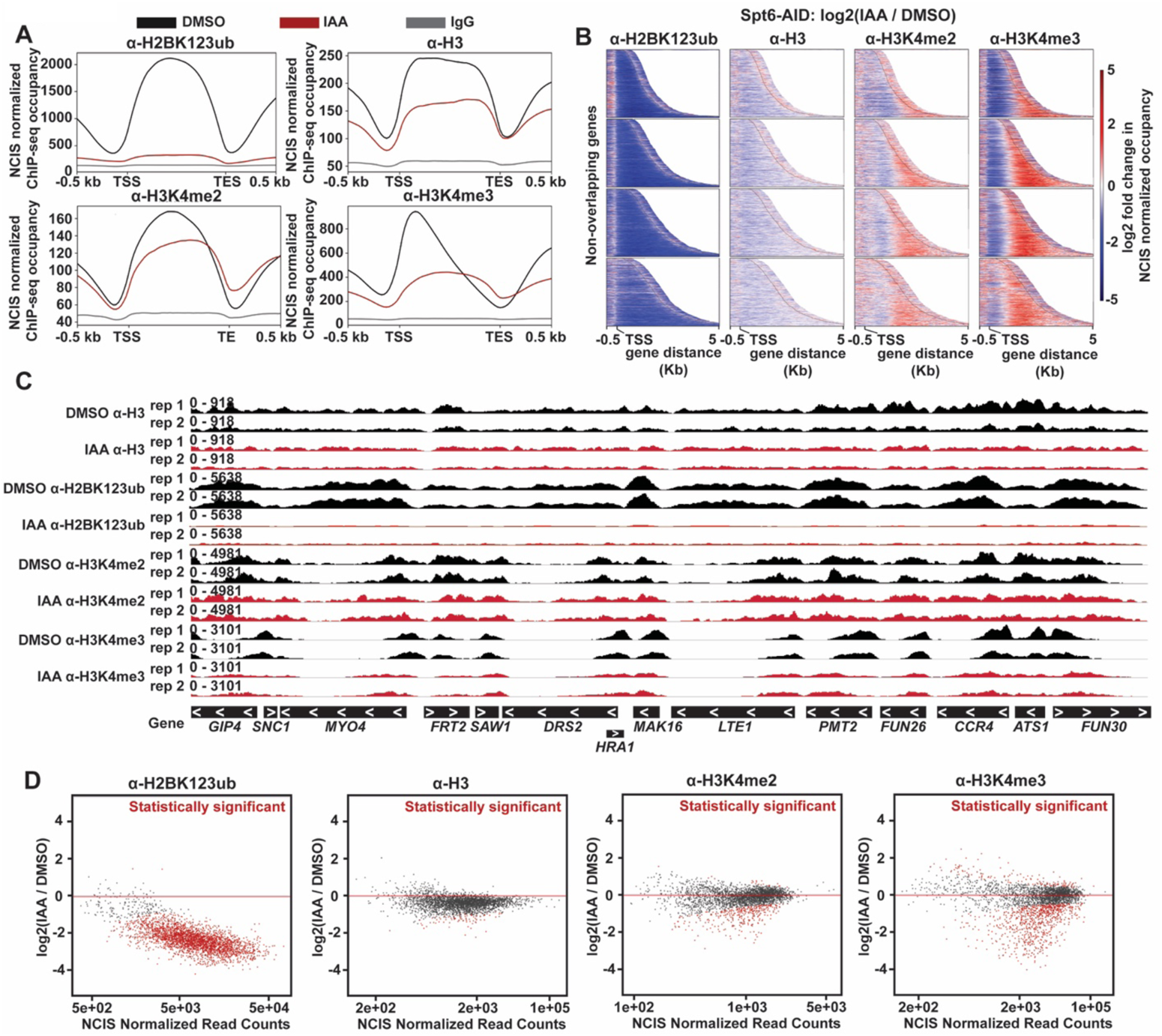
Acute depletion of Spt6 leads to changes in Paf1C-dependent histone modifications genome-wide. (**A**) Aggregation plots of NCIS-normalized ChIP-seq data for H2BK123ub, total H3, H3K4me2, and H3K4me3. IgG was used as a background control for ChIP-seq. (**B**) Heatmaps of differential H2BK123ub, total H3, H3K4me2, and H3K4me3 occupancy (IAA vs DMSO) with genes divided into quartiles based on Rpb3 occupancy and sorted by length. Heatmaps show the region between the transcription start site (TSS) and the cleavage and poly-adenylation site (labeled with a black line) +/- 500 bp and use a bin size of 25 bp. (**C**) Genome browser tracks of a region of chromosome 1 (chrI:84,656-118,640) showing NCIS-normalized ChIP-seq occupancy for individual biological replicates. (**D**) MA plots showing differential occupancy, determined by DESeq2 of NCIS-normalized ChIP-seq data (IAA vs DMSO), for H2BK123ub, total H3, H3K4me2, and H3K4me3. Genes with statistically significant changes in occupancy (FDR adjusted p value ≤ 0.05) are colored red. (**A, B, D**) The results presented are derived from 3087 non-overlapping protein-coding genes (92) and were generated by averaging data from two biological replicates.

## DISCUSSION

In this study, we investigated the recruitment of Paf1C to the active Pol II elongation complex, focusing on the role of the Cdc73 subunit. Early studies in yeast revealed that null mutations in *CDC73* and *RTF1* dissociated Paf1C from chromatin and pointed to these two subunits as important mediators of the interaction of Paf1C with Pol II (93). The interaction between Rtf1, through its Plus3 domain, and the phosphorylated Spt5 CTR is understood at the structural level, and mutations that alter this interaction reduce Paf1C occupancy on chromatin in yeast (32,39,40). However, the interactions between Cdc73 and the Pol II elongation complex had not been fully defined, due in part to limited structural information on Cdc73 within the elongation complex (4, 5). We previously showed that the Cdc73 C-domain, which adopts a Ras-like fold, is required for full Paf1C occupancy on transcribed genes but not Paf1C assembly (42). Here we identify Spt6 and Rpb1 as direct interactors of Cdc73 *in vivo*, show that Cdc73 and Spt6 can interact *in vitro* in the absence of other factors, and demonstrate a requirement for Spt6 in recruiting Paf1C to chromatin genome-wide.

Prior to this work, evidence suggested an important role for the phosphorylated Pol II CTD in recruiting Paf1C to chromatin. In both yeast and mammalian cells, inactivation of protein kinases that phosphorylate the CTD lower Paf1C occupancy on active genes or association of Paf1C with isolated elongation complexes (31,33,34,94). These kinases include Kin28/CDK8, which phosphorylates serines 5 and 7 of the CTD heptapeptide repeat, and Bur1/CDK9 and CDK12, which phosphorylate serine 2 of the repeat [reviewed in (95)]. Peptide pulldown assays with purified yeast Paf1C or isolated subunits provided more direct evidence for an interaction between Paf1C and the phosphorylated Pol II CTD. These studies detected binding of three different Paf1C subunits, Cdc73, Rtf1, and Ctr9, to CTD repeat peptides phosphorylated at positions Ser2 and Ser5 or Ser5 and Ser7 (33). The requirement for phosphorylation at serines 2, 5, and 7 for these *in vitro* interactions is consistent with the specificities of the kinases implicated by cell-based assays to be important for Paf1C recruitment. The peptide interacting region of Cdc73 mapped to amino acids 201-393, which overlap the structurally defined C-domain (amino acids 230-393) (42), and substitution of hydrophobic residues within the C-domain reduced Cdc73 occupancy at several active genes (33). While these studies support an interaction between Cdc73 and the phosphorylated CTD, interpretations are complicated by the ability of three different subunits to bind the same CTD peptides and the ability of these same subunits to bind to phosphorylated Spt5 peptides (33). Our BPA crosslinking data provide the first evidence for a direct interaction between the Cdc73 C-domain and Rpb1 *in vivo*. Using a protease-cleavable form of Rpb1, we mapped this interaction to the last 271 amino acids of Rpb1, which contains both the CTD and a recently defined Rpb1 linker region that is phosphorylated by Bur1 and interacts with the Spt6 tSH2 domain (79, 96). From the Cdc73 side of the interaction, substitution of W321 to alanine greatly diminished the Cdc73-Rpb1 crosslink. Among a large number of amino acid substitutions tested in our previous mutational analysis of the C-domain, the Cdc73 W321A substitution was the only one to confer detectable sensitivity to the base analog 6-azauracil, a property shared by other elongation factor mutants (42). Compared to relevant single mutant strains, *cdc73-W321A rtf1Δ* double mutant strains exhibit enhanced sensitivity to 6-azauracil and lower levels of H3K36me3, consistent with the idea that Cdc73 and Rtf1 separably impact Paf1C recruitment and function (42).

Our search for Cdc73 interactors by *in vivo* BPA crosslinking also identified an interaction between Cdc73 and Spt6. Spt6 is an essential, conserved member of the Pol II elongation complex whose occupancy on chromatin correlates with that of Pol II (3,4,37,38). Spt6 promotes transcription elongation (17,87,88,97), interacts with histones and a co-regulatory protein Spn1 (98–101), stimulates Set2-mediated H3K36me3 (102–104), and functions as a histone chaperone to maintain chromatin integrity on active genes (86,105,106). To our knowledge, our results are the first demonstration of a direct interaction between Cdc73 and Spt6, with the exception of a small number of chemically induced crosslinks between Spt6 and Cdc73 within the reconstituted human active elongation complex (4). Surprisingly, the same BPA-substituted residue in Cdc73, R300, crosslinked to Rpb1 and Spt6, and both crosslinks were reduced by the W321A substitution. These results might reflect physical proximity between all three proteins within the active elongation complex, dynamic changes in the conformation of this complex, or subpopulations of elongation complexes.

Using purified recombinant proteins, we demonstrated that an Spt6 derivative lacking the acidic, unstructured N-terminal region (65) is sufficient to interact with Cdc73 *in vitro*. Through chemical crosslinking and structural modeling, we identified a putative interface between the Cdc73 C-domain and the YqgF, HhH, and DLD domains of Spt6. An Spt6 derivative lacking the YqgF, HhH, and DLD domains was completely defective for binding to Cdc73 *in vitro*. In addition, an Spt6 derivative lacking the S1 domain and residues linking it to the YqgF domain was greatly reduced for Cdc73 binding, suggesting that the S1 domain is important for the Cdc73 interaction either directly or possibly indirectly through stabilizing the Spt6 core. We tested the importance of the predicted Cdc73-Spt6 interface by targeted mutational analysis but failed to observe a significant effect on the chromatin occupancy of Paf1C, even in a sensitized background where the Rtf1-Spt5 interaction has been mutationally compromised. It is possible that the interface predicted by our crosslinking and modeling studies of isolated proteins does not play a major role in Paf1C recruitment or that redundancy in the molecular interface allows minor perturbations to be tolerated.

To place it within the context of the Pol II elongation complex, we aligned our model of the Cdc73-Spt6 interaction to the cryo-EM structure of the complete, activated Pol II elongation complex (EC*), which contains Pol II, PAF (Paf1, Ctr9, Cdc73, Leo1, Rtf1 and Wdr61), Spt6, and DSIF (5). Only a portion of hCdc73, corresponding to residues 218-260, is observed in this EC* structure. This region, termed the anchor helices/μ-catenin binding region, interacts with Ctr9. Residues 218-260 in the human protein correspond to 159-168 in the yeast protein (based on a ClustalO alignment). Alignment of our model of the proposed Cdc73-Spt6 interface with the complete EC* structure revealed a distance of ∼162 Å between the C-terminus of the hCdc73 anchor helices and the N-terminus of the yCdc73 C-domain. The 62 amino acids between residue 168 and 230 in the yeast protein (at Cα to Cα of 3.5 Å) allow for a maximum distance of 217 Å, suggesting that placement of the C-domain as predicted by its interaction with Spt6 is compatible with the location of the anchor helices in the complete EC*. This would place the C-domain near the Spt5 KOWx-KOW4 domains and the stalk of Pol II near the nascent RNA. Validation of the Cdc73-Spt6 interaction requires further structural studies.

Our biochemical results led us to investigate a role for Spt6 in Paf1C recruitment and/or maintenance on chromatin. Previous studies showed that Paf1C occupancy on chromatin was diminished by the *S. cerevisiae spt6-1004* mutation, the analogous mutation in *Schizosaccharomyces pombe SPT6*, or depletion of Drosophila Spt6 by RNAi (47,89,97). These ChIP results are supported by a recent proteomic study, which showed a significant reduction in Pol II-associated Paf1C in strains containing the *spt6-1004* mutation, an in-frame deletion that removes the Spt6 HhH domain (80). While these studies indicate a role for Spt6, and possibly the HhH domain more directly, in controlling Paf1C occupancy on chromatin, interpretations are complicated by the instability of the Spt6-1004 protein and of some Paf1C subunits in *spt6-1004* mutant strains (47, 80). We chose two alternative approaches to test the requirement for Spt6 in recruitment of Paf1C to Pol II. First, using the *spt6-50* mutation, we dissociated Spt6 from Pol II and observed a dramatic decrease in Paf1C occupancy on active genes. Second, to rule out indirect effects caused by long-term absence of a functional Spt6 protein, which might explain the reduction in Spt5 occupancy in the *spt6-50* background, we used the AID system to acutely deplete Spt6 and observed genome-wide loss of Paf1 occupancy. Unlike Paf1, Spt5 occupancy on chromatin was not decreased upon rapid Spt6 depletion, suggesting that Spt5 does not require Spt6 for its recruitment to Pol II. Under these conditions, effects on Paf1 occupancy cannot be easily explained by loss of the Rtf1-Spt5 interaction, although we cannot rule out conformational changes to the Rtf1-Spt5 interface in the absence of Spt6.

Consistent with findings by Jeronimo *et al*. (90), we found that loss of Spt6 broadly affects histone modification patterns genome-wide, including those that are Paf1C-dependent. Acute depletion of Spt6 caused a large reduction in H2BK123ub on gene bodies, which can be explained by the inability to reestablish this modification in the absence of Paf1C after pre-existing H2Bub marks are turned over by the deubiquitylases Ubp8 and Ubp10 (107). As noted previously (90), we also observed a redistribution of H3K4me2 and H3K4me3 toward the 3’ ends of genes in the absence of Spt6. The retention of H3 K4me2 and H3 K4me3, albeit in a mislocalized pattern, is consistent with the stable nature of these modifications in yeast (108). Therefore, through multiple activities, which include recruitment of Paf1C, recycling of histones during transcription elongation (90), and association with the histone chaperone Spn1 (92), Spt6 plays a major role in determining histone modification patterns genome-wide.

In conclusion, through the identification of interactions between Cdc73 and two components of the core Pol II elongation machinery, our findings provide new insights into the coupling of Paf1C to Pol II, a process that is important for the patterning of Paf1C-mediated histone modifications and the control of transcription elongation. Interestingly, three mediators of Paf1C attachment to Pol II --- the Spt5 CTR, the Rpb1 CTD, and the Rpb1 linker region, which binds to Spt6 --- are all targets of the Bur1/CDK9 kinase, providing an explanation for the strong requirement for this kinase in governing Paf1C recruitment (31, 33). Understanding why multiple interactions within the Pol II elongation complex are required for Paf1C recruitment and how these interactions are regulated will be important areas for future study. Finally, we note that the Cdc73-Spt6 and Cdc73-Rpb1 interactions identified here offer a Paf1C recruitment mechanism independent of Rtf1, which might prove instrumental in human cells where hRtf1 is loosely associated with hPaf1C.

## DATA AVAILABILITY

Strains and plasmids are available upon request. ChIP-seq data have been deposited in the Gene Expression Omnibus database under accession number GSE201436. The code used for the analysis of ChIP-seq data has been uploaded to the following GitHub repository: https://github.com/mae92/Spt6_Project_ChIPseq_Analysis. Information related to modeling of the Cdc73-Spt6 interaction, including PDB files, scripts, and crosslinking data, can be accessed at the following GitHub repository: https://github.com/mae92/Spt6_Project_XLMS_Structural_Modeling.

## SUPPLEMENTARY DATA

Supplementary Data have been submitted with the manuscript.

## ACKNOWLEDGMENTS

We thank Roni Lahr and Andrea Berman for advice on quantifying protein binding assays, Andrew VanDemark and members of his laboratory for sharing equipment and providing technical advice particularly on protein interaction assays, Craig Kaplan for the *RPB1*-TEV plasmid, Fred Winston and Nathan Clark for yeast strains, Tim Formosa, Grant Hartzog and John Woolford for antibodies, and Julia Seraly for assisting with the analysis of the *spt6* mutants. We are grateful for the helpful suggestions we have received from Frank Pugh, Olga Viktorovskaya, and Fred Winston. We also thank Craig Kaplan, Sarah Hainer, Miler Lee, members of their laboratories, and all members of the Arndt laboratory, especially Alex Francette, for many helpful discussions. This project used the University of Pittsburgh Health Sciences Sequencing Core at UPMC Children’s Hospital of Pittsburgh for genomic sequencing experiments and was supported in part by the University of Pittsburgh Center for Research Computing through the resources provided.

