## supplemental figures and tables for "Spt6 directly interacts with Cdc73 and is required for Paf1C recruitment to active genes"

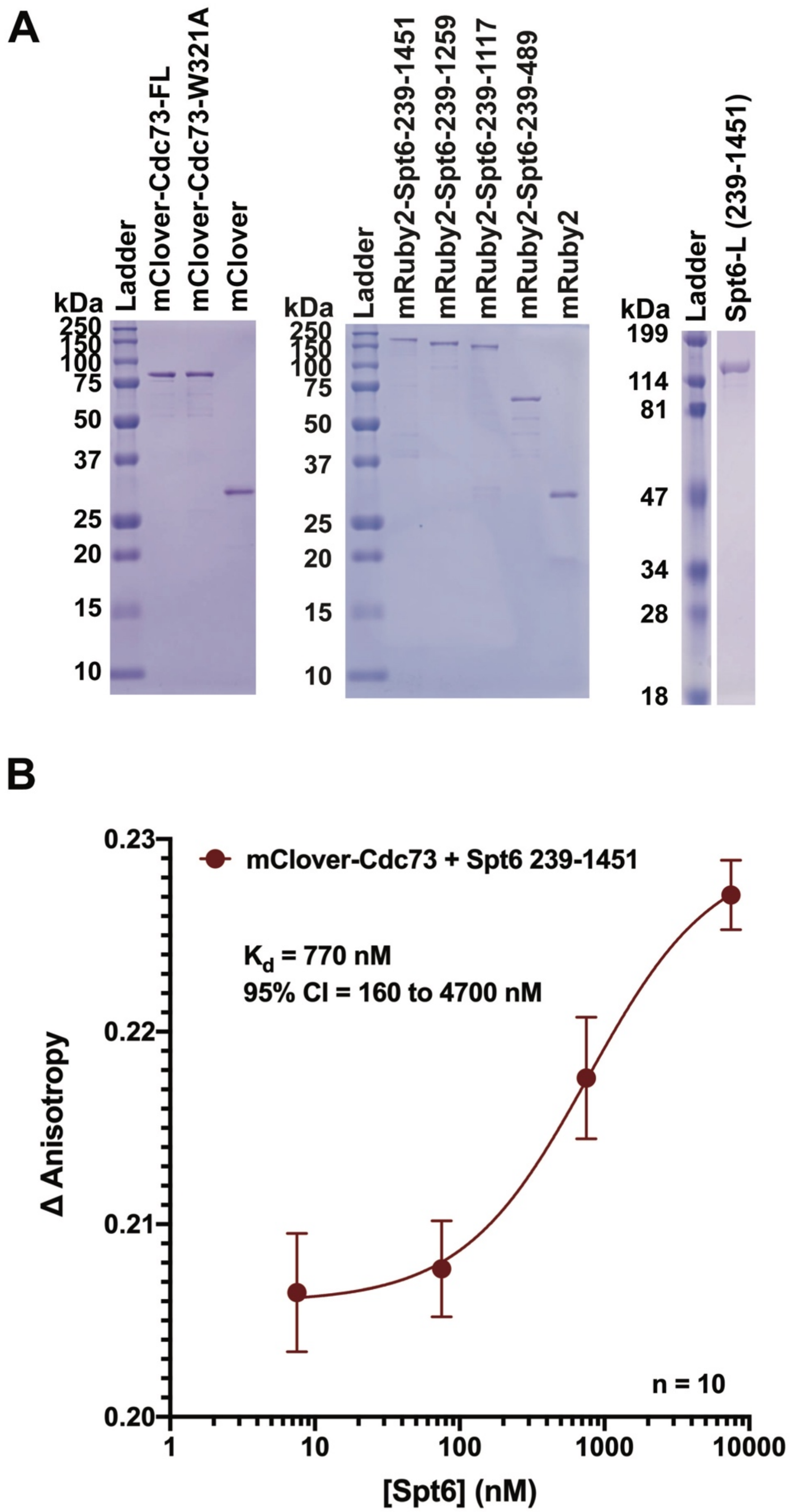
A
kDa
Ladder
mClover-Cdc73-FL
mClover-Cdc73-W321A
mClover
250
150
100
75
50
37
25
20
15
10
kDa
Ladder
mRuby2-Spt6-239-1451
mRuby2-Spt6-239-1259
mRuby2-Spt6-239-1117
mRuby2-Spt6-239-489
mRuby2
250
150
100
75
50
37
25
20
15
10
kDa
Ladder
Spt6-L (239-1451)
199
114
81
47
34
28
18
B
mClover-Cdc73 + Spt6 239-1451
K_d = 770 nM
95% CI = 160 to 4700 nM
Δ Anisotropy
0.23
0.22
0.21
0.20
n = 10
1
10
100
1000
10000
[Spt6] (nM)

**Figure S1. Protein expression and fluorescence anisotropy binding assay results.** (**A**) SDS PAGE gel stained with Coomassie blue to show purity of indicated recombinant proteins. (**B**) Fluorescence anisotropy analysis of Spt6 (239-1451) binding to full-length 10xHis-mClover-Cdc73. The data represent the average of ten experiments. Binding curves were fit to data using Prism 8 software as described in the Methods.

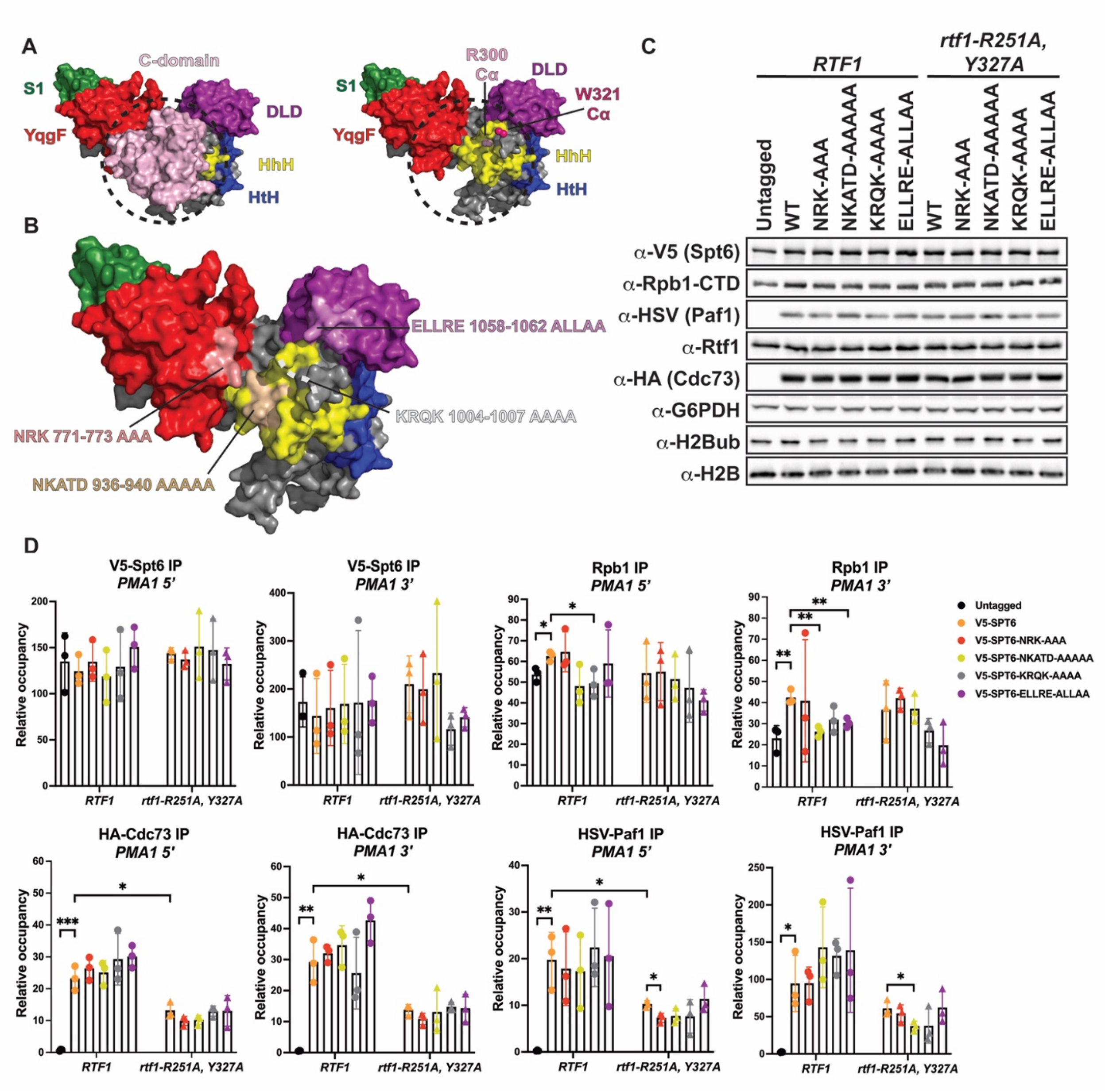

**Figure S2. Mutational analysis of the putative binding interface between Cdc73 and Spt6**. (**A**) Left: Structural model of the Spt6-Cdc73 C-domain interaction. Right: Locations of the Cdc73 residues mutated in Figure 1 are indicated on the putative interface. For visualization of these residues, the bulk of Cdc73 has been removed from this image. Spt6 domains are colored as in Figure 2B. (**B**) Spt6 alanine scanning substitutions targeted to the putative Spt6-Cdc73 interface. (**C**) Western analysis of *spt6* mutants assessing protein levels of V5-Spt6, Rpb1 (8WG16), Paf1C subunits and H2B K123ub (H2Bub). Wild-type *V5-SPT6* (WT) and V5-tagged *spt6* mutations, encoding the amino acid substitutions shown in B, were introduced by plasmid shuffling into *RTF1*

(KY3698 and KY3699) and *rtf1-R251A, Y327A* (KY3917 and KY3918) genetic backgrounds. G6PDH and H2B serve as loading controls. Representative gels from three biological replicates are shown. Untagged indicates a strain with untagged Paf1C components and WT V5-Spt6. (**D**) ChIP-qPCR analysis of V5-Spt6, Rpb1-CTD (8WG16), HA-Cdc73 and HSV-Paf1 occupancy at the 5’ and 3’ regions of a highly expressed gene, *PMA1*. Data are normalized to input. The mean and standard deviation of three biological replicates are plotted. Unpaired student’s t test *$p < 0.05$; **$p < 0.01$; ***$p < 0.001$.

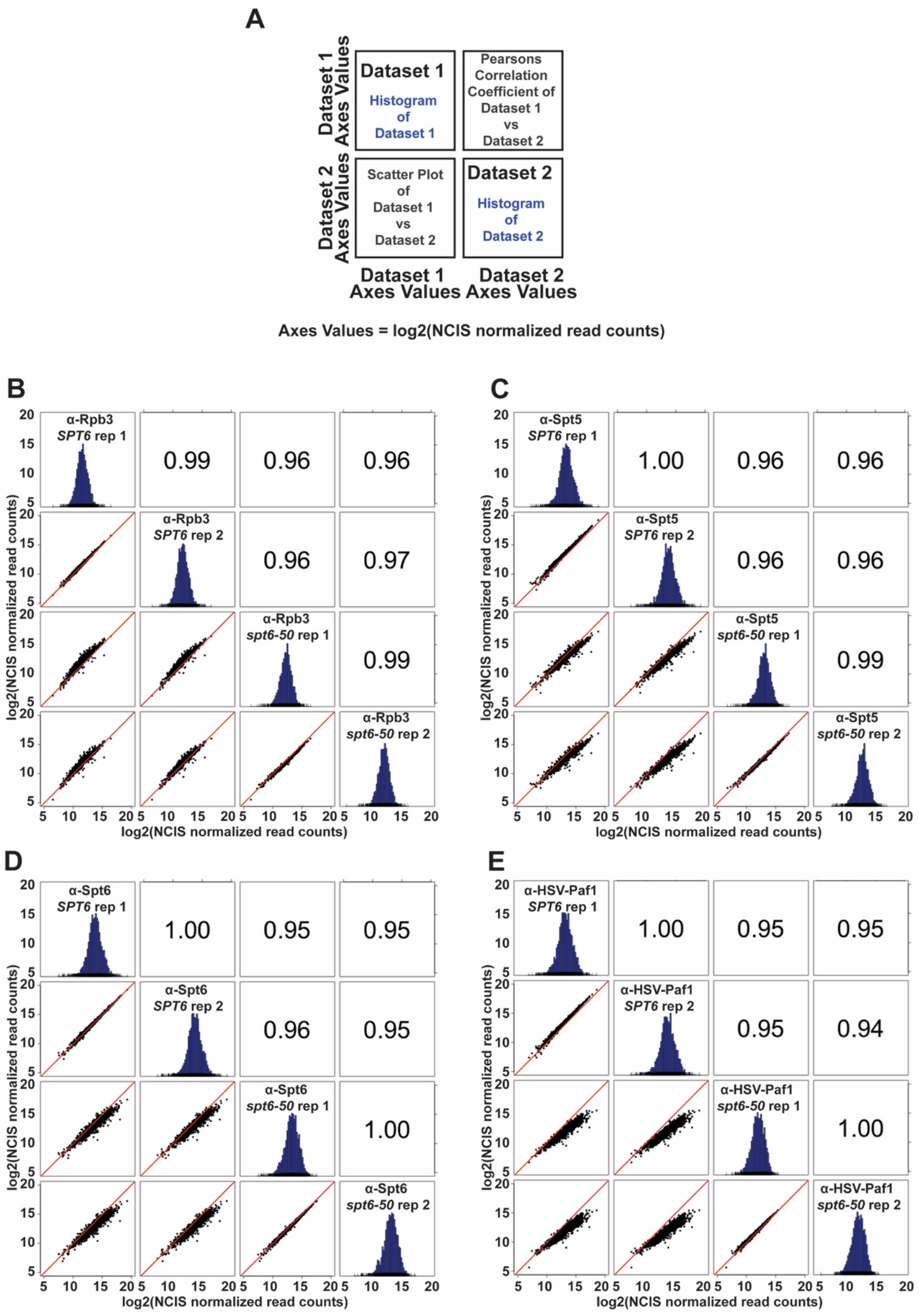
A
Dataset 1
Histogram of Dataset 1
Pearsons Correlation Coefficient of Dataset 1 vs Dataset 2
Scatter Plot of Dataset 1 vs Dataset 2
Dataset 2
Histogram of Dataset 2
Dataset 1 Axes Values
Dataset 2 Axes Values
Axes Values = log2(NCIS normalized read counts)
B
α-Rpb3 SPT6 rep 1
α-Rpb3 SPT6 rep 2
α-Rpb3 spt6-50 rep 1
α-Rpb3 spt6-50 rep 2
0.99
0.96
0.96
0.96
0.97
0.99
log2(NCIS normalized read counts)
C
α-Spt5 SPT6 rep 1
α-Spt5 SPT6 rep 2
α-Spt5 spt6-50 rep 1
α-Spt5 spt6-50 rep 2
1.00
0.96
0.96
0.96
0.96
0.99
D
α-Spt6 SPT6 rep 1
α-Spt6 SPT6 rep 2
α-Spt6 spt6-50 rep 1
α-Spt6 spt6-50 rep 2
1.00
0.95
0.95
0.96
0.95
1.00
E
α-HSV-Paf1 SPT6 rep 1
α-HSV-Paf1 SPT6 rep 2
α-HSV-Paf1 spt6-50 rep 1
α-HSV-Paf1 spt6-50 rep 2
1.00
0.95
0.95
0.95
0.94
1.00

**Figure S3. Correlation analysis of *SPT6* and *spt6-50* ChIP-seq data.** (**A**) Diagram illustrating how the correlation data are plotted. (**B, C, D, E**) Scatter plots, histograms, and Pearson correlation coefficients of NCIS-normalized ChIP-seq data for Rpb3 (**B**), Spt5 (**C**), Spt6 (**D**), and HSV-Paf1 (**E**). The red line represents x = y. (**B, C, D, E**) The results presented are from 3087 non-overlapping protein-coding genes (92).

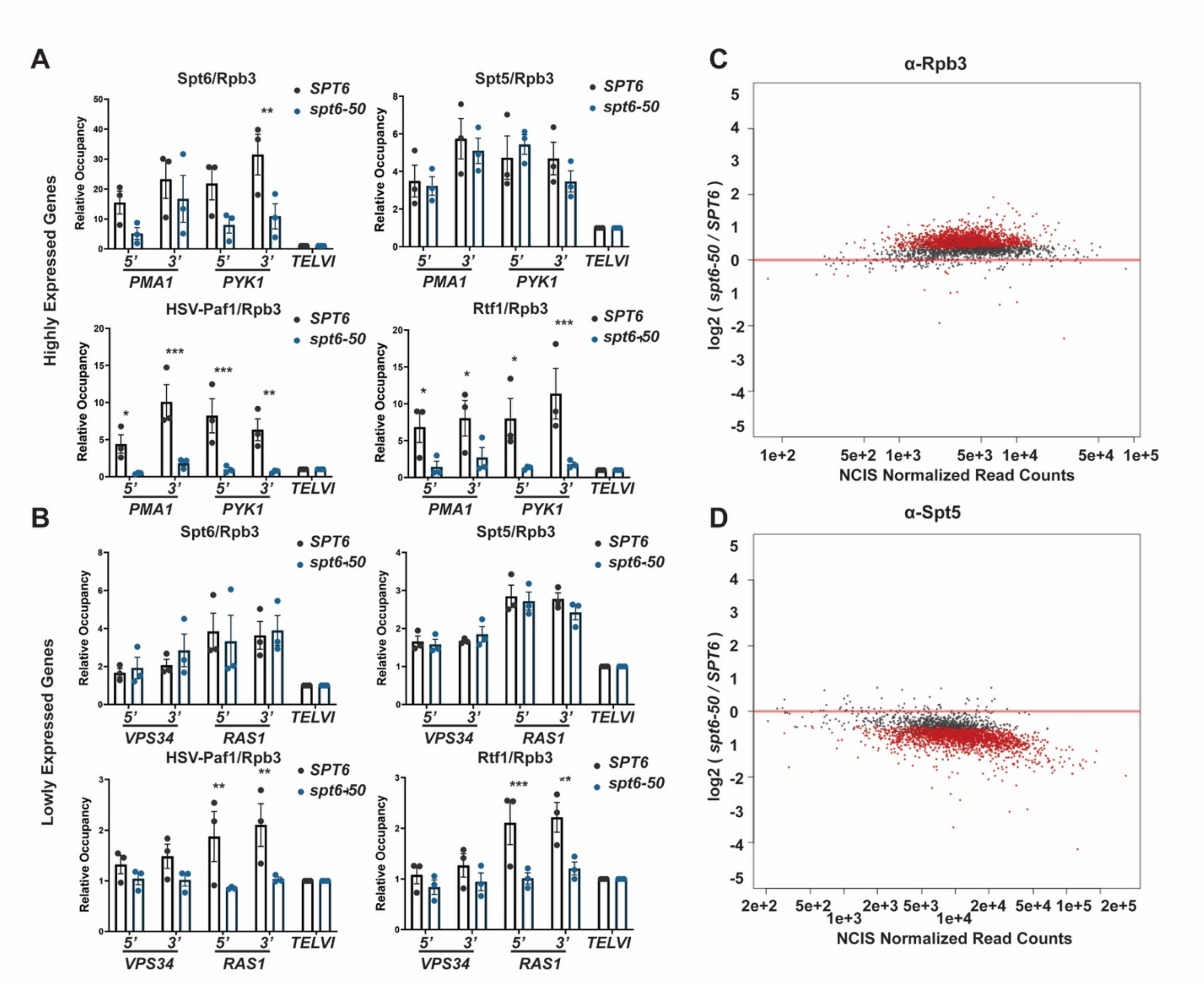

**Figure S4. Chromatin occupancy of elongation factors in *SPT6* and *spt6-50* strains analyzed by ChIP-qPCR and ChIP-seq.** (**A, B**) ChIP-qPCR analysis of Spt6, Spt5, HSV-Paf1, and Rtf1 occupancy plotted relative to Rpb3 occupancy for two highly expressed genes, *PMA1* and *PYK1* (**A**), and two lowly expressed genes, *VPS34* and *RAS1* (**B**). Highly and lowly expressed genes were identified from Rpb3 ChIP-seq data with highly expressed genes having 5- 20 fold higher occupancy than lowly expressed genes. ChIP-qPCR data for target genes were normalized both to input and ChIP signals for a region near telomere VI. (**A, B**) The mean and standard error of three biological replicates are plotted. Unpaired student's t test *$p < 0.05$; **$p < 0.01$; ***$p < 0.001$. (**C, D**) MA plots showing differential occupancy for Rpb3 and Spt5 determined by DESeq2 analysis of NCIS-normalized ChIP-seq data (*spt6-50* vs *SPT6*). Genes with statistically significant changes in occupancy (FDR-adjusted p value $\leq$ 0.05) are colored red.

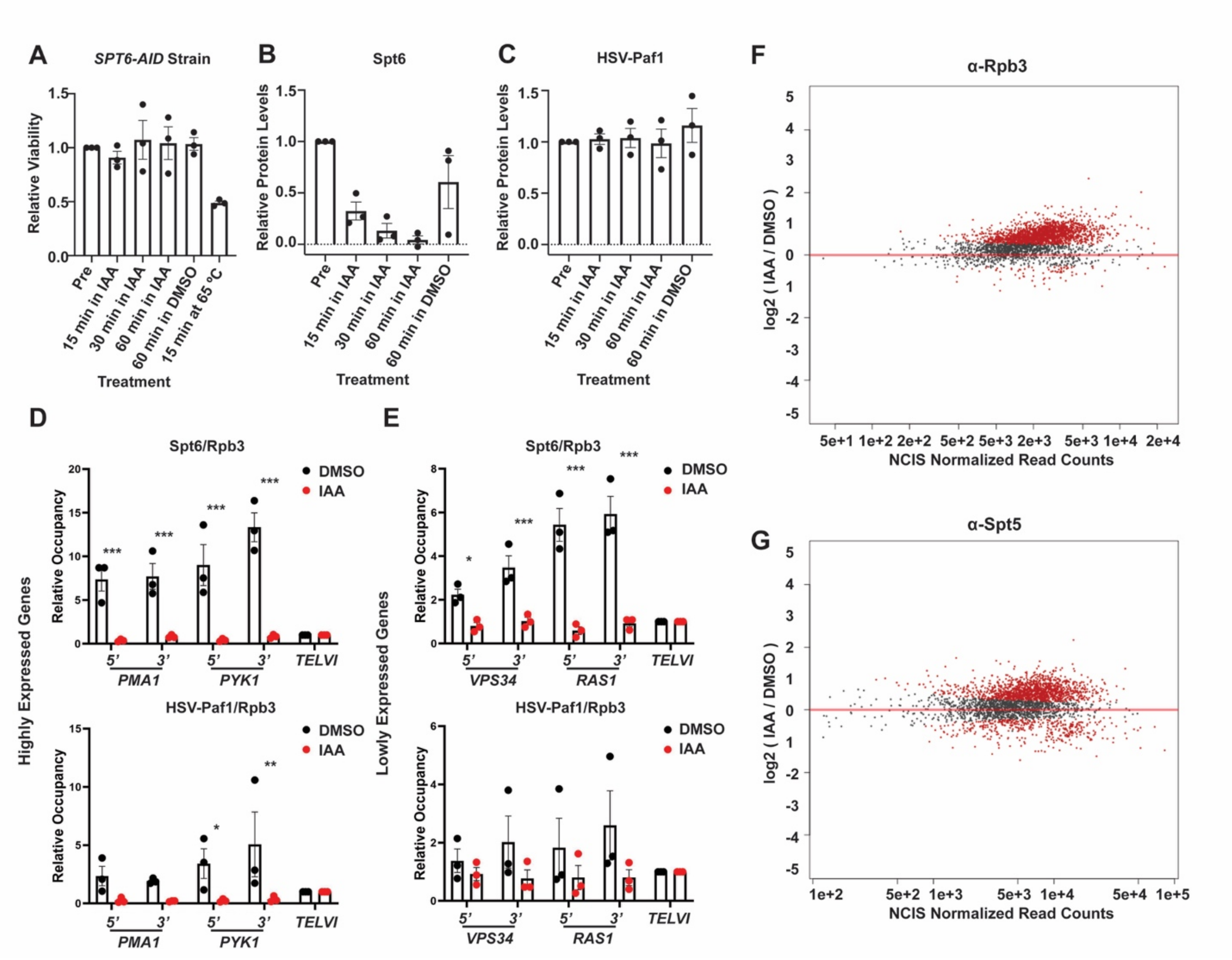

**Figure S5. Chromatin occupancy of Spt6 and Paf1 in *SPT6* and *SPT6-AID* strains analyzed by ChIP-qPCR and ChIP-seq.** (**A**) Relative cell viability determined by methylene blue staining of *SPT6-AID* cells following treatment with IAA or DMSO and sampling at 15, 30, and 60 min (IAA), 60 min (DMSO) or 60 min at 65 ˚C (as a control). Viability was calculated relative to the pre-treatment sample. Three biological replicates were analyzed. (**B, C**) Quantification of protein levels for three biological replicates assayed by western blot (representative blot is shown in Fig. 5B). Spt6 and HSV-Paf1 signals were normalized to the loading control (G6PDH) and the ratio for the pretreatment sample was set to 1. (**D, E**) ChIP-qPCR analysis of Spt6 and HSV-Paf1 plotted relative to Rpb3 occupancy for two highly expressed genes, *PMA1* and *PYK1* (**D**), and two lowly expressed genes, *VPS34* and *RAS1* (**E**), as defined in the legend to Supplementary Figure 4. ChIP-qPCR data for target genes were normalized both to input and ChIP signals for a region near telomere VI. (**A, B**) The mean and standard error of three biological replicates are plotted. Unpaired student's t test **p* < 0.05; ***p* < 0.01; ****p* < 0.001. (**F, G**) MA plots showing differential occupancy for Rpb3 and Spt5, determined by DESeq2 analysis of NCIS-normalized ChIP-seq data (IAA vs DMSO). Genes with statistically significant changes in occupancy (FDR-adjusted p value < 0.05)

are colored red. (**F, G**) The results presented are from 3087 non-overlapping protein-coding genes (92).

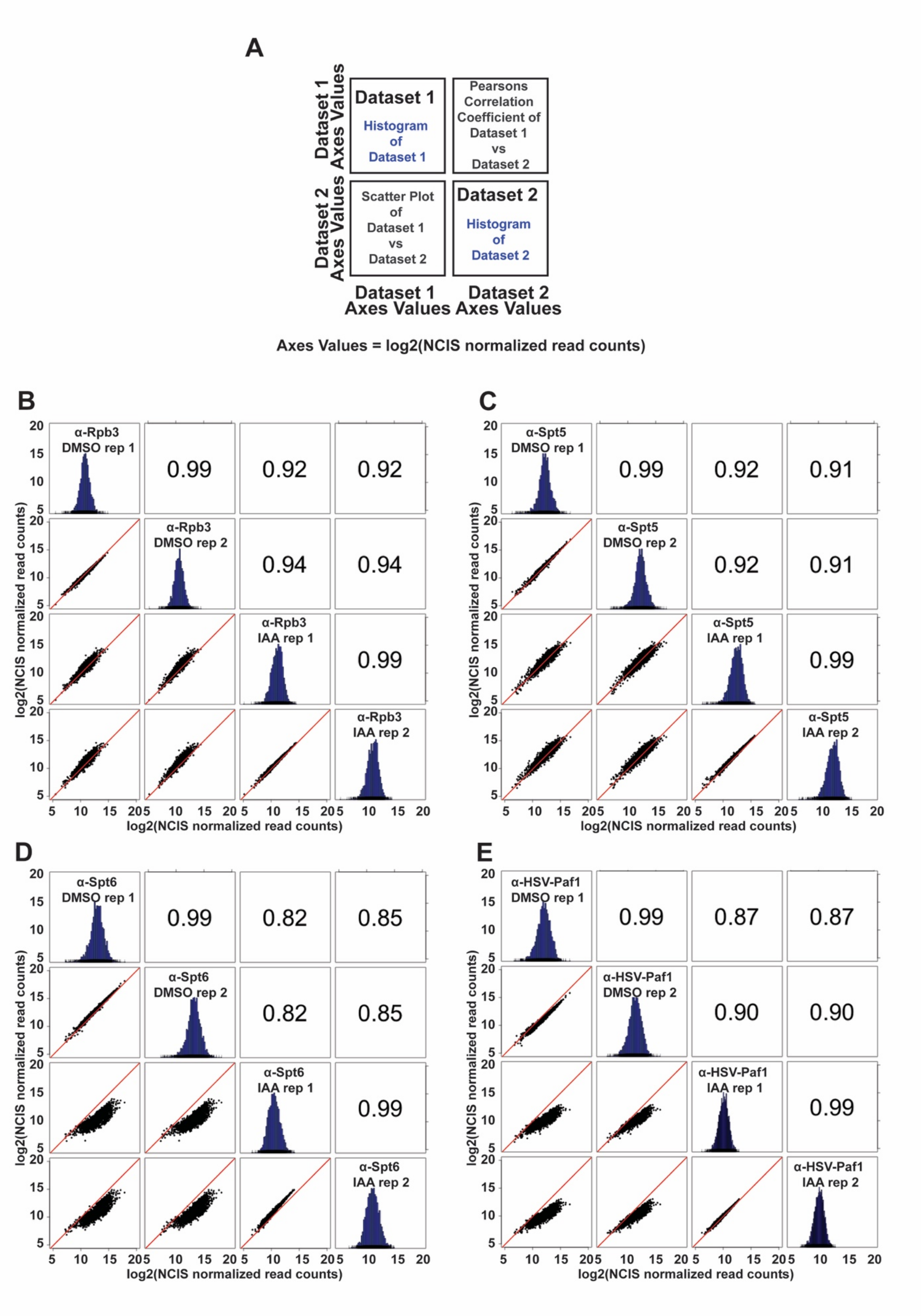
A
Dataset 1
Histogram of Dataset 1
Pearsons Correlation Coefficient of Dataset 1 vs Dataset 2
Scatter Plot of Dataset 1 vs Dataset 2
Dataset 2
Histogram of Dataset 2
Dataset 1 Axes Values
Dataset 2 Axes Values
Axes Values = log2(NCIS normalized read counts)
B
α-Rpb3 DMSO rep 1
α-Rpb3 DMSO rep 2
α-Rpb3 IAA rep 1
α-Rpb3 IAA rep 2
0.99
0.92
0.92
0.94
0.94
0.99
C
α-Spt5 DMSO rep 1
α-Spt5 DMSO rep 2
α-Spt5 IAA rep 1
α-Spt5 IAA rep 2
0.99
0.92
0.91
0.92
0.91
0.99
D
α-Spt6 DMSO rep 1
α-Spt6 DMSO rep 2
α-Spt6 IAA rep 1
α-Spt6 IAA rep 2
0.99
0.82
0.85
0.82
0.85
0.99
E
α-HSV-Paf1 DMSO rep 1
α-HSV-Paf1 DMSO rep 2
α-HSV-Paf1 IAA rep 1
α-HSV-Paf1 IAA rep 2
0.99
0.87
0.87
0.90
0.90
0.99
log2(NCIS normalized read counts)
5
10
15
20

**Figure S6. Correlation analysis of Rpb3 and transcription elongation factor ChIP-seq data obtained with the *SPT6-AID* strain treated with DMSO or IAA.** (**A**) Diagram illustrating how the correlation data are plotted. (**B, C, D, E**) Scatter plots, histograms, and Pearson correlation coefficients of NCIS-normalized ChIP-seq data for Rpb3 (**B**), Spt5 (**C**), Spt6 (**D**), HSV-Paf1 (**E**). The red line represents x = y. (**B, C, D, E**) The results presented are from 3087 non-overlapping protein-coding genes (92).

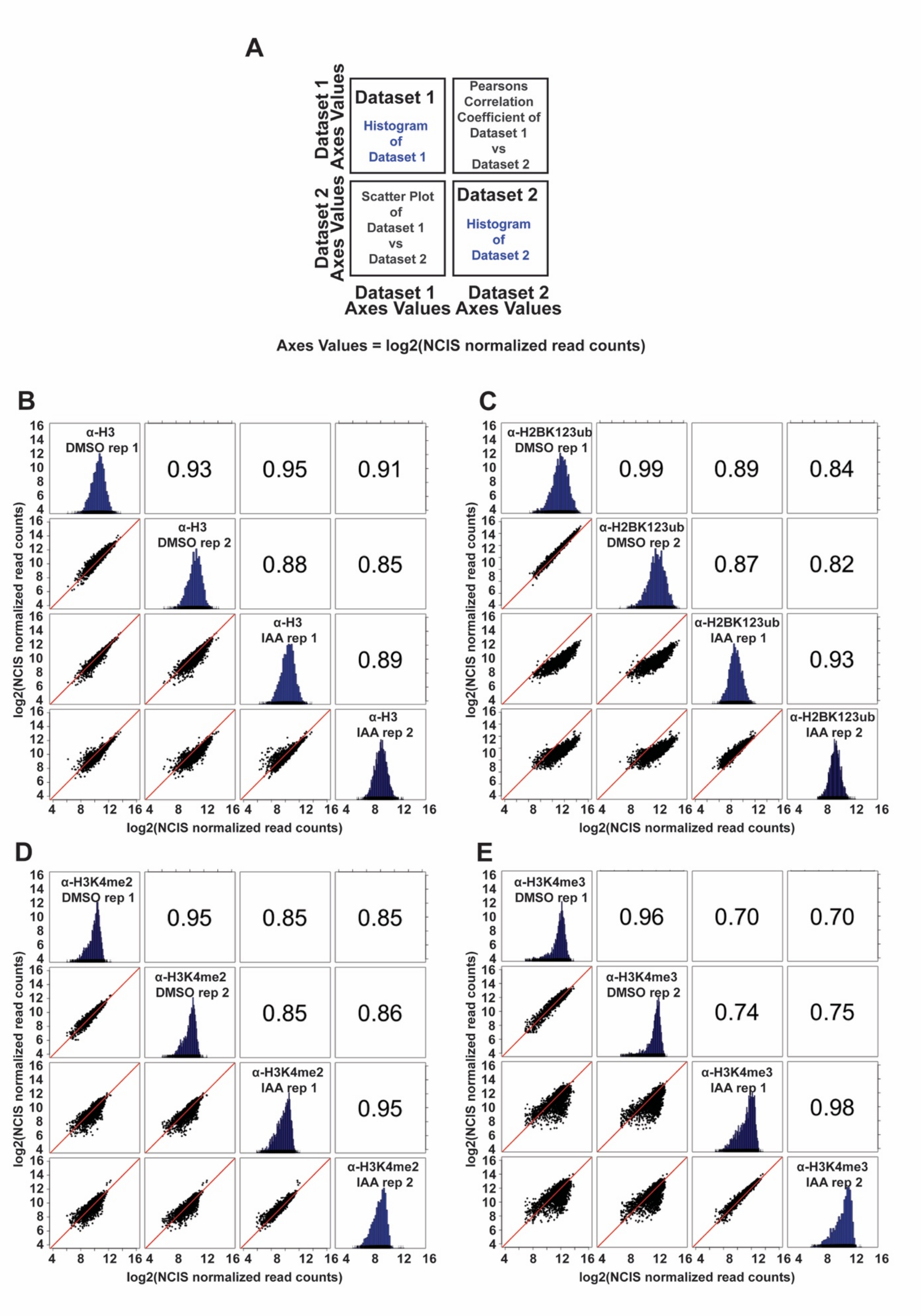

A
Dataset 1
Histogram of Dataset 1
Pearsons Correlation Coefficient of Dataset 1 vs Dataset 2
Scatter Plot of Dataset 1 vs Dataset 2
Dataset 2
Histogram of Dataset 2
Dataset 1 Axes Values
Dataset 2 Axes Values
Axes Values = log2(NCIS normalized read counts)
B
α-H3 DMSO rep 1
α-H3 DMSO rep 2
α-H3 IAA rep 1
α-H3 IAA rep 2
0.93
0.95
0.91
0.88
0.85
0.89
C
α-H2BK123ub DMSO rep 1
α-H2BK123ub DMSO rep 2
α-H2BK123ub IAA rep 1
α-H2BK123ub IAA rep 2
0.99
0.89
0.84
0.87
0.82
0.93
D
α-H3K4me2 DMSO rep 1
α-H3K4me2 DMSO rep 2
α-H3K4me2 IAA rep 1
α-H3K4me2 IAA rep 2
0.95
0.85
0.85
0.85
0.86
0.95
E
α-H3K4me3 DMSO rep 1
α-H3K4me3 DMSO rep 2
α-H3K4me3 IAA rep 1
α-H3K4me3 IAA rep 2
0.96
0.70
0.70
0.74
0.75
0.98
log2(NCIS normalized read counts)

**Figure S7. Correlation analysis of histone and histone modification ChIP-seq data obtained with the *SPT6-AID* strain treated with DMSO or IAA.** (**A**) Diagram illustrating how the correlation data are plotted. (**B, C, D, E**) Scatter plots, histograms, and Pearson correlation coefficients of NCIS-normalized ChIP-seq data for total H3 (**B**), H2BK123ub (**C**), H3K4me2 (**D**), H3K4me3 (**E**). The red line represents x = y. (**B, C, D, E**) The results are from 3087 non-overlapping protein-coding genes (92).

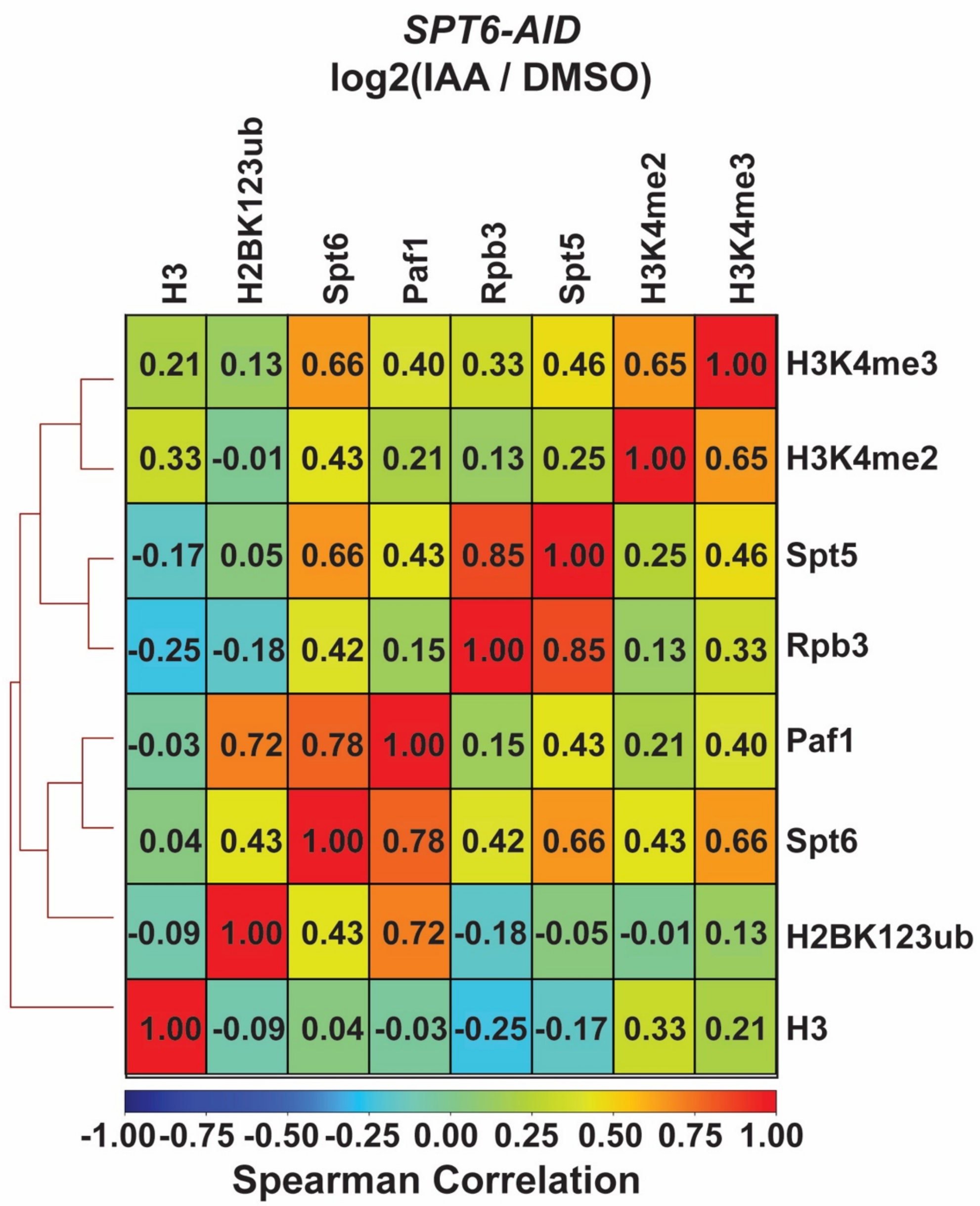

**Figure S8. Comparison of ChIP-seq data for elongation factor and histone modification occupancies in *SPT6-AID* samples.** Spearman correlation coefficients were calculated from the average of two biological replicates per ChIP-seq experiment.

### Supplementary Table S1. Yeast strains

| Strain | Genotype |
|---|---|
| BTY42 | *MAT**a** his3Δ200 leu2Δ1 trp1Δ63 cdc73Δ::KANMX* |
| KY303 | *MATα his4-912∂ lys2-128∂ leu2Δ1 ura3-52* |
| KY600 | *MAT**a** his4-912∂ lys2-128∂ leu2Δ1 spt6-14* |
| KY1203 | *MATα his3Δ200 leu2Δ1 ura3-52 trp1Δ63 arg4-12 cdc73Δ::KanMX rpb1Δ187::HIS3 [pRP112=RPB1 URA3 CEN/ARS]* |
| KY1855 | *MATα his4-912∂ leu2Δ1 ura3-52 trp1Δ63 cdc73Δ::KANMX* |
| KY1858 | *MAT**a** his4-912∂ trp1Δ63 cdc73Δ::KANMX* |
| KY2242 | *MATα his4-912∂ lys2-128∂ trp1Δ63 cdc73Δ::KANMX* |
| KY2838 | *MATα his3Δ200 leu2Δ1 ura3-52 trp1Δ63 arg4-12 cdc73Δ::KANMX rpb1Δ187::HIS3 [pCKB858 = rpb1::1461TEV, LEU2, CEN/ARS]* |
| KY3208 | *MAT**a** his3Δ200::HIS3::osTIR lys2-128∂ leu2Δ1 ura3-52* |
| KY3291 | *MAT**a** his3Δ200::HIS3-osTIR lys2-128∂ leu2Δ1 ura3-52 trp1Δ63 3XHSV-PAF1 SPT6-V5-IAA7(AID)-KANMX* |
| KY3343 | *MAT**a** his3Δ200::HIS3-osTIR lys2-128∂ leu2Δ1 ura3-52 3XHSV-PAF1* |
| KY3352 | *MAT**a** his3Δ200::HIS3-osTIR lys2-173R2 leu2Δ1 ura3-52 trp1Δ63 ade8 spt6-50 3XHSV-PAF1* |
| KY3638 | *MATα lys2-128∂ leu2Δ1 trp1Δ63 ade8 3XHSV-PAF1 CDC73-HA::kanMX spt6-1004* |
| KY3642 | *MATα leu2Δ1 trp1Δ63 3XHSV-PAF1 CDC73-HA::kanMX* |
| KY3698 | *MATα his4-912∂ lys2-128∂ leu2Δ1 ura3-52 trp1Δ63 spt6Δ::LEU2 [pCC11 = SPT6, URA3, CEN/ARS]* |
| KY3699 | *MATα his4-912∂ lys2-128∂ leu2Δ1 ura3-52 trp1Δ63 spt6Δ::LEU2 3XHSV-PAF1 CDC73-HA::kanMX [pCC11 = SPT6, URA3, CEN/ARS]* |

| | |
|---|---|
| KY3900 | *MATα his4-912∂ lys2-128∂ leu2Δ1 ura3-52 trp1Δ63 spt6Δ::LEU2 3XHSV-PAF1 CDC73-HA::kanMX [pKB1623 = SPT6-3XV5, TRP1]* |
| KY3907 | *MATα his4-912∂ lys2-128∂ leu2Δ1 ura3-52 trp1Δ63 spt6Δ::LEU2 3XHSV-PAF1 CDC73-HA::kanMX rtf1Δ236-331:: pCORE-Hyg,Ura [pKB1623 = 3XV5-SPT6, TRP1,CEN/ARS]* |
| KY3908 | *MATα his4-912∂ lys2-128∂ leu2Δ1 ura3-52 trp1Δ63 spt6Δ::LEU2 3XHSV-PAF1 CDC73-HA::kanMX rtf1Δ236-331:: pCORE-Hyg,Ura [pKB1623 = 3XV5-SPT6, TRP1,CEN/ARS]* |
| KY3915 | *MATα his4-912∂ lys2-128∂ leu2Δ1 ura3-52 trp1Δ63 spt6Δ::LEU2 3XHSV-PAF1 CDC73-HA::kanMX rtf1- R351A,Y327A [pKB1623 = 3XV5-SPT6, TRP1,CEN/ARS]* |
| KY3916 | *MATα his4-912∂ lys2-128∂ leu2Δ1 ura3-52 trp1Δ63 spt6Δ::LEU2 3XHSV-PAF1 CDC73-HA::kanMX rtf1- R351A,Y327A [pKB1623 = 3XV5-SPT6, TRP1,CEN/ARS]* |
| KY3917 | *MATα his4-912∂ lys2-128∂ leu2Δ1 ura3-52 trp1Δ63 spt6Δ::LEU2 3XHSV-PAF1 CDC73-HA::kanMX rtf1- R351A,Y327A [pCK25 = SPT6-FLAG, URA3, CEN/ARS]* |
| KY3918 | *MATα his4-912∂ lys2-128∂ leu2Δ1 ura3-52 trp1Δ63 spt6Δ::LEU2 3XHSV-PAF1 CDC73-HA::kanMX rtf1- R351A,Y327A [pCK25 = SPT6-FLAG, URA3, CEN/ARS]* |
| KL01 | *K. lactis* spike-in (prototroph) |
| KP07 | *S. pombe h- ura4-D18 ade6::ade6+ spt5::spt5+-3xV5-IAA17-KanMX6 Padh15-skp1-OsTIR1-natMX6-Padh15-skp1-AtTIR1-2NLS-9myc* |
| KP08 | *S. pombe h+ ura4-D18 leu1-32 ade6-m210 spt5+::3HA-KanMX hrp1::HSV-hphMX6* |

### Supplementary Table S2. Plasmids

| Name | Alias | Insert Description | *E. coli* Marker | *S. cerevisiae* Marker | Type | Source | Ref. |
|---|---|---|---|---|---|---|---|
| pLH157 | pKB1027 | *ADH1$_p$-EC Tyr* RS | Amp$^R$ | *TRP1* | 2µ | Steve Hahn | (109) |
| pEK01 | pKB1273 | *ADH1$_p$ 3XHSV-CDC73* | Kan$^R$ | *TRP1* | 2µ | this paper | |
| pEK02 | pKB1274 | *ADH1$_p$ 3XHSV-cdc73 W321Amber* | Kan$^R$ | *TRP1* | 2µ | this paper | |
| pEK03 | pKB1275 | *ADH1$_p$ 3XHSV-cdc73 H319Amber* | Kan$^R$ | *TRP1* | 2µ | this paper | |
| pEK04 | pKB1276 | *ADH1$_p$ 3XHSV-cdc73 R268Amber* | Kan$^R$ | *TRP1* | 2µ | this paper | |
| pEK05 | pKB1277 | *ADH1$_p$ 3XHSV-cdc73 S272Amber* | Kan$^R$ | *TRP1* | 2µ | this paper | |
| pEK06 | pKB1278 | *ADH1$_p$ 3XHSV-cdc73 R300Amber* | Kan$^R$ | *TRP1* | 2µ | this paper | |
| pCK858 | pKB1341 | *rpb1::1461TEV* | Amp$^R$ | *LEU2* | *CEN/ARS* | Craig Kaplan | |
| | pKB1347 | *ADH1$_p$ 3XHSV-cdc73 W321A* | Kan$^R$ | *TRP1* | 2µ | this paper | |
| | pKB1349 | *ADH1$_p$ 3XHSV-cdc73 W321A R300Amber* | Kan$^R$ | *TRP1* | 2µ | this paper | |
| | pKB1359 | *ADH1$_p$ 3XHSV-CDC73* | Kan$^R$ | *URA3* | 2µ | this paper | |
| | pKB1361 | *ADH1$_p$ 3XHSV-cdc73 R268Amber* | Kan$^R$ | *URA3* | 2µ | this paper | |
| pSB2065 | pKB1435 | 3XV5-IAA7-KanMX | Amp$^R$ | *KanMX* | *integrating* | Sue Biggins | (110) |
| pSB2273 | pKB1436 | *GPD1-osTIR-HIS3* | Amp$^R$ | *HIS3* | *integrating* | Sue Biggins | (110) |
| pJZ1 | | 10xHis-mRuby2 in pET28a | Kan$^R$ | | *expression* | Andrew VanDemark | (111) |
| pJZ2 | | 10xHis-mClover in pET28a | Kan$^R$ | | *expression* | Andrew VanDemark | (111) |
| | pKB1453 | 10xHis-mRuby2-Spt6-239-1451 | Kan$^R$ | | *expression* | this paper | |
| | pKB1454 | 10xHis-mRuby2-Spt6-239-1259 | Kan$^R$ | | *expression* | this paper | |
| | pKB1482 | 10xHis-mClover-Cdc73 | Kan$^R$ | | *expression* | this paper | |
| | pKB1504 | 10xHis-mClover-Cdc73-W321A | Kan$^R$ | | *expression* | this paper | |
| | pKB1506 | 10xHis-mRuby2-Spt6-239-1117 | Kan$^R$ | | *expression* | this paper | |
| | pKB1507 | 10xHis-mRuby2-Spt6-239-489 | Kan$^R$ | | *expression* | this paper | |
| | pKB1573 | Cdc73-10xHis | Amp$^R$ | | *expression* | this paper | |
| pCK30 + V5 | pKB1623 | *3XV5-SPT6* | Amp$^R$ | *TRP1* | *CEN/ARS* | Craig Kaplan | |
| | pKB1636 | *3XV5-spt6-NRK771-773AAA* | Amp$^R$ | *TRP1* | *CEN/ARS* | this paper | |

| | | | | | | |
|---|---|---|---|---|---|---|
| | pKB1638 | *3XV5-spt6-NKATD936-940AAAAA* | $Amp^R$ | *TRP1* | *CEN/ARS* | this paper |
| | pKB1641 | *3XV5-spt6-KRQK1004-1007AAAA* | $Amp^R$ | *TRP1* | *CEN/ARS* | this paper |
| | pKB1645 | *3XV5-spt6-ELLRE1058-1062ALLAA* | $Amp^R$ | *TRP1* | *CEN/ARS* | this paper |

### Supplementary Table S3. Primers used for qPCR experiments

| Primer | Target | Sequence (5' to 3') | Efficiency |
|---|---|---|---|
| APO95 | *TEL VI* FWD | TGCAAGCGTAACAAAGCCATA | 2.04 |
| APO96 | *TEL VI* REV | TCCGAACGCTATTCCAGAAAG | |
| ECO234 | *PMA1* 5' FWD | GCTAGACCAGTTCCAGAAGAATATTTACA | 1.97 |
| ECO235 | *PMA1* 5' REV | CAGCCATTTGATTCAAACCGTA | |
| ECO236 | *PMA1* 3' FWD | GAAATCTTCTTGGGTCTATGGATTG | 1.94 |
| ECO237 | *PMA1* 3' REV | CAACATCAGCGAAAATAGCGAT | |
| ECO238 | *PYK1* 5' FWD | ACCAAGGGTCCAGAAATCAGAAC | 1.98 |
| ECO239 | *PYK1* 5' REV | TGTCATCGGTGGTGAAGATCAT | |
| ECO240 | *PYK1* 3' FWD | AGAAACTGTACTCCAAAGCCAACCT | 1.99 |
| ECO241 | *PYK1* 3' REV | TGGTCTGTACTTGGAAACCAATCTT | |
| MEO185 | *YPS34* 3' FWD | CTTCGAACATACCTGATATCAGAATAG | 1.90 |
| MEO186 | *YPS34* 3' REV | GCACTGTGGCATCTTCTTCGGAC | |
| MEO183 | *YPS34* 5' FWD | GTGTCTCACAGGATCTGGATGTTCC | 1.91 |
| MEO184 | *YPS34* 5' REV | GAGATGGCTTCAACAGTGGCTTATG | |
| MEO189 | *RAS1* 3' FWD | CGAGAAGTAAACAGTCTGCTGAG | 1.92 |
| MEO190 | *RAS1* 3' REV | CAAATTATACAACAACCACCACTAG | |
| MEO187 | *RAS1* 5' FWD | GCAGGGAAATAAATCAACTATAAGAG | 2.00 |
| MEO188 | *RAS1* 5' REV | GATAGTAGGGTCATATTCGTCCAC | |

**Supplementary Table S4. All DSS and EDC crosslink sites for mClover-Cdc73 and Spt6 (239-1451).** Inter-protein crosslinks used in IMP modeling are highlighted in bold.

| Experiments with Crosslink Observed | Total Experiments | Crosslinker | Protein 1 | AA 1 | Protein 2 | AA 2 |
|---|---|---|---|---|---|---|
| 1 | 2 | EDC | Cdc73 | 24 | Cdc73 | 205 |
| 1 | 2 | EDC | Cdc73 | 24 | Cdc73 | 371 |
| 2 | 2 | EDC | Cdc73 | 119 | Cdc73 | 126 |
| 1 | 2 | EDC | Cdc73 | 119 | Cdc73 | 141 |
| 2 | 2 | EDC | Cdc73 | 130 | Cdc73 | 141 |
| 2 | 2 | EDC | Cdc73 | 156 | Cdc73 | 205 |
| 1 | 2 | EDC | Cdc73 | 158 | Cdc73 | 205 |
| 2 | 2 | EDC | Cdc73 | 176 | Cdc73 | 205 |
| 1 | 2 | EDC | Cdc73 | 176 | Cdc73 | 371 |
| 2 | 2 | EDC | Cdc73 | 185 | Cdc73 | 176 |
| 1 | 2 | EDC | Cdc73 | 205 | Cdc73 | 141 |
| 1 | 2 | EDC | Cdc73 | 263 | Cdc73 | 24 |
| **2** | **2** | **EDC** | **Cdc73** | **24** | **Spt6** | **834** |
| **1** | **2** | **EDC** | **Cdc73** | **185** | **Spt6** | **940** |
| **2** | **2** | **EDC** | **Cdc73** | **185** | **Spt6** | **1062** |
| **2** | **2** | **EDC** | **Cdc73** | **185** | **Spt6** | **1063** |
| **1** | **2** | **EDC** | **Cdc73** | **194** | **Spt6** | **1062** |
| **1** | **2** | **EDC** | **Cdc73** | **199** | **Spt6** | **1062** |
| **1** | **2** | **EDC** | **Cdc73** | **205** | **Spt6** | **971** |
| **1** | **2** | **EDC** | **Cdc73** | **205** | **Spt6** | **1009** |
| **2** | **2** | **EDC** | **Cdc73** | **205** | **Spt6** | **1012** |
| **1** | **2** | **EDC** | **Cdc73** | **205** | **Spt6** | **1038** |
| **1** | **2** | **EDC** | **Cdc73** | **205** | **Spt6** | **1048** |
| **1** | **2** | **EDC** | **Cdc73** | **205** | **Spt6** | **1049** |
| **2** | **2** | **EDC** | **Cdc73** | **205** | **Spt6** | **1063** |
| **1** | **2** | **EDC** | **Cdc73** | **205** | **Spt6** | **1071** |
| **2** | **2** | **EDC** | **Cdc73** | **205** | **Spt6** | **1076** |
| **2** | **2** | **EDC** | **Cdc73** | **304** | **Spt6** | **353** |
| **1** | **2** | **EDC** | **Cdc73** | **304** | **Spt6** | **940** |
| **1** | **2** | **EDC** | **Cdc73** | **304** | **Spt6** | **1012** |
| **2** | **2** | **EDC** | **Cdc73** | **304** | **Spt6** | **1062** |
| **2** | **2** | **EDC** | **Cdc73** | **371** | **Spt6** | **345** |
| **1** | **2** | **EDC** | **Cdc73** | **371** | **Spt6** | **940** |

| | | | | | | |
|---|---|---|---|---|---|---|
| 1 | 2 | EDC | Spt6 | 301 | Spt6 | 307 |
| 1 | 2 | EDC | Spt6 | 306 | Spt6 | 312 |
| 1 | 2 | EDC | Spt6 | 306 | Spt6 | 861 |
| 1 | 2 | EDC | Spt6 | 314 | Spt6 | 319 |
| 1 | 2 | EDC | Spt6 | 335 | Spt6 | 306 |
| 1 | 2 | EDC | Spt6 | 335 | Spt6 | 383 |
| 1 | 2 | EDC | Spt6 | 342 | Spt6 | 431 |
| 1 | 2 | EDC | Spt6 | 384 | Spt6 | 379 |
| 2 | 2 | EDC | Spt6 | 462 | Spt6 | 431 |
| 1 | 2 | EDC | Spt6 | 699 | Spt6 | 1239 |
| 2 | 2 | EDC | Spt6 | 940 | Spt6 | 773 |
| 2 | 2 | EDC | Spt6 | 1007 | Spt6 | 971 |
| 1 | 2 | EDC | Spt6 | 1007 | Spt6 | 1055 |
| 1 | 2 | EDC | Spt6 | 1007 | Spt6 | 1058 |
| 2 | 2 | EDC | Spt6 | 1007 | Spt6 | 1062 |
| 2 | 2 | EDC | Spt6 | 1007 | Spt6 | 1063 |
| 1 | 2 | EDC | Spt6 | 1009 | Spt6 | 773 |
| 2 | 2 | EDC | Spt6 | 1069 | Spt6 | 1038 |
| 2 | 2 | EDC | Spt6 | 1184 | Spt6 | 1278 |
| 1 | 2 | EDC | Spt6 | 1230 | Spt6 | 319 |
| 1 | 2 | EDC | Spt6 | 1232 | Spt6 | 611 |
| 2 | 2 | EDC | Spt6 | 1232 | Spt6 | 1239 |
| 2 | 2 | EDC | Spt6 | 1235 | Spt6 | 1239 |
| 2 | 2 | EDC | Spt6 | 1236 | Spt6 | 611 |
| 1 | 2 | EDC | Spt6 | 1237 | Spt6 | 611 |
| 2 | 3 | DSS | Cdc73 | 77 | Cdc73 | 17 |
| 3 | 3 | DSS | Cdc73 | 119 | Cdc73 | 130 |
| 2 | 3 | DSS | Cdc73 | 119 | Cdc73 | 136 |
| 2 | 3 | DSS | Cdc73 | 119 | Cdc73 | 205 |
| 1 | 3 | DSS | Cdc73 | 119 | Cdc73 | 304 |
| 2 | 3 | DSS | Cdc73 | 119 | Cdc73 | 371 |
| 1 | 3 | DSS | Cdc73 | 124 | Cdc73 | 205 |
| 2 | 3 | DSS | Cdc73 | 124 | Cdc73 | 304 |
| 1 | 3 | DSS | Cdc73 | 124 | Cdc73 | 371 |
| 3 | 3 | DSS | Cdc73 | 130 | Cdc73 | 185 |
| 1 | 3 | DSS | Cdc73 | 130 | Cdc73 | 205 |
| 3 | 3 | DSS | Cdc73 | 130 | Cdc73 | 371 |
| 1 | 3 | DSS | Cdc73 | 146 | Cdc73 | 130 |
| 2 | 3 | DSS | Cdc73 | 169 | Cdc73 | 185 |

| | | | | | | |
|---|---|---|---|---|---|---|
| 2 | 3 | DSS | Cdc73 | 169 | Cdc73 | 205 |
| 2 | 3 | DSS | Cdc73 | 169 | Cdc73 | 304 |
| 3 | 3 | DSS | Cdc73 | 169 | Cdc73 | 371 |
| 1 | 3 | DSS | Cdc73 | 185 | Cdc73 | 199 |
| 3 | 3 | DSS | Cdc73 | 185 | Cdc73 | 205 |
| 2 | 3 | DSS | Cdc73 | 185 | Cdc73 | 371 |
| 3 | 3 | DSS | Cdc73 | 194 | Cdc73 | 205 |
| 1 | 3 | DSS | Cdc73 | 205 | Cdc73 | 17 |
| 3 | 3 | DSS | Cdc73 | 205 | Cdc73 | 212 |
| 3 | 3 | DSS | Cdc73 | 205 | Cdc73 | 371 |
| 1 | 3 | DSS | Cdc73 | 218 | Cdc73 | 205 |
| 2 | 3 | DSS | Cdc73 | 226 | Cdc73 | 205 |
| 1 | 3 | DSS | Cdc73 | 236 | Cdc73 | 205 |
| 3 | 3 | DSS | Cdc73 | 236 | Cdc73 | 226 |
| 3 | 3 | DSS | Cdc73 | 236 | Cdc73 | 304 |
| 1 | 3 | DSS | Cdc73 | 263 | Cdc73 | 366 |
| 3 | 3 | DSS | Cdc73 | 263 | Cdc73 | 371 |
| 2 | 3 | DSS | Cdc73 | 282 | Cdc73 | 205 |
| 3 | 3 | DSS | Cdc73 | 282 | Cdc73 | 371 |
| 3 | 3 | DSS | Cdc73 | 304 | Cdc73 | 17 |
| 3 | 3 | DSS | Cdc73 | 304 | Cdc73 | 130 |
| 2 | 3 | DSS | Cdc73 | 304 | Cdc73 | 185 |
| 3 | 3 | DSS | Cdc73 | 304 | Cdc73 | 205 |
| 1 | 3 | DSS | Cdc73 | 304 | Cdc73 | 212 |
| 2 | 3 | DSS | Cdc73 | 304 | Cdc73 | 218 |
| 3 | 3 | DSS | Cdc73 | 304 | Cdc73 | 226 |
| 3 | 3 | DSS | Cdc73 | 304 | Cdc73 | 371 |
| 1 | 3 | DSS | Cdc73 | 361 | Cdc73 | 371 |
| 3 | 3 | DSS | Cdc73 | 366 | Cdc73 | 371 |
| 1 | 3 | DSS | Cdc73 | 368 | Cdc73 | 366 |
| 3 | 3 | DSS | Cdc73 | 368 | Cdc73 | 371 |
| 3 | 3 | DSS | Cdc73 | 371 | Cdc73 | 366 |
| 2 | 3 | DSS | Cdc73 | 371 | Cdc73 | 368 |
| 1 | 3 | DSS | Cdc73 | 385 | Cdc73 | 218 |
| 2 | 3 | DSS | Cdc73 | 385 | Cdc73 | 226 |
| 1 | 3 | DSS | Cdc73 | 385 | Cdc73 | 304 |
| 3 | 3 | DSS | Cdc73 | 385 | Cdc73 | 371 |
| **1** | **3** | **DSS** | **Cdc73** | **185** | **Spt6** | **316** |
| **2** | **3** | **DSS** | **Cdc73** | **205** | **Spt6** | **319** |

| | | | | | | |
|---|---|---|---|---|---|---|
| **3** | **3** | **DSS** | **Cdc73** | **263** | **Spt6** | **1134** |
| **1** | **3** | **DSS** | **Cdc73** | **304** | **Spt6** | **319** |
| **1** | **3** | **DSS** | **Cdc73** | **304** | **Spt6** | **728** |
| **1** | **3** | **DSS** | **Cdc73** | **304** | **Spt6** | **956** |
| **1** | **3** | **DSS** | **Cdc73** | **304** | **Spt6** | **1134** |
| **1** | **3** | **DSS** | **Cdc73** | **304** | **Spt6** | **1239** |
| **1** | **3** | **DSS** | **Cdc73** | **366** | **Spt6** | **1239** |
| **3** | **3** | **DSS** | **Cdc73** | **371** | **Spt6** | **319** |
| **1** | **3** | **DSS** | **Cdc73** | **371** | **Spt6** | **616** |
| **1** | **3** | **DSS** | **Cdc73** | **371** | **Spt6** | **728** |
| **3** | **3** | **DSS** | **Cdc73** | **371** | **Spt6** | **870** |
| **1** | **3** | **DSS** | **Cdc73** | **371** | **Spt6** | **1007** |
| **1** | **3** | **DSS** | **Cdc73** | **371** | **Spt6** | **1134** |
| **1** | **3** | **DSS** | **Cdc73** | **371** | **Spt6** | **1239** |
| **1** | **3** | **DSS** | **Cdc73** | **385** | **Spt6** | **319** |
| **1** | **3** | **DSS** | **Cdc73** | **385** | **Spt6** | **1239** |
| 2 | 3 | DSS | Spt6 | 306 | Spt6 | 316 |
| 2 | 3 | DSS | Spt6 | 306 | Spt6 | 319 |
| 2 | 3 | DSS | Spt6 | 306 | Spt6 | 715 |
| 2 | 3 | DSS | Spt6 | 306 | Spt6 | 956 |
| 3 | 3 | DSS | Spt6 | 306 | Spt6 | 1357 |
| 3 | 3 | DSS | Spt6 | 307 | Spt6 | 319 |
| 3 | 3 | DSS | Spt6 | 307 | Spt6 | 956 |
| 3 | 3 | DSS | Spt6 | 316 | Spt6 | 319 |
| 3 | 3 | DSS | Spt6 | 316 | Spt6 | 956 |
| 1 | 3 | DSS | Spt6 | 316 | Spt6 | 958 |
| 2 | 3 | DSS | Spt6 | 354 | Spt6 | 715 |
| 3 | 3 | DSS | Spt6 | 354 | Spt6 | 719 |
| 3 | 3 | DSS | Spt6 | 354 | Spt6 | 723 |
| 1 | 3 | DSS | Spt6 | 379 | Spt6 | 319 |
| 1 | 3 | DSS | Spt6 | 383 | Spt6 | 319 |
| 1 | 3 | DSS | Spt6 | 504 | Spt6 | 1444 |
| 2 | 3 | DSS | Spt6 | 576 | Spt6 | 1444 |
| 2 | 3 | DSS | Spt6 | 598 | Spt6 | 354 |
| 1 | 3 | DSS | Spt6 | 598 | Spt6 | 591 |
| 2 | 3 | DSS | Spt6 | 598 | Spt6 | 719 |
| 1 | 3 | DSS | Spt6 | 609 | Spt6 | 1239 |
| 2 | 3 | DSS | Spt6 | 616 | Spt6 | 1134 |
| 3 | 3 | DSS | Spt6 | 616 | Spt6 | 1239 |

| | | | | | | |
|---|---|---|---|---|---|---|
| 1 | 3 | DSS | Spt6 | 700 | Spt6 | 576 |
| 1 | 3 | DSS | Spt6 | 700 | Spt6 | 584 |
| 3 | 3 | DSS | Spt6 | 728 | Spt6 | 723 |
| 3 | 3 | DSS | Spt6 | 746 | Spt6 | 773 |
| 3 | 3 | DSS | Spt6 | 749 | Spt6 | 773 |
| 3 | 3 | DSS | Spt6 | 773 | Spt6 | 746 |
| 1 | 3 | DSS | Spt6 | 834 | Spt6 | 773 |
| 3 | 3 | DSS | Spt6 | 870 | Spt6 | 781 |
| 1 | 3 | DSS | Spt6 | 870 | Spt6 | 834 |
| 1 | 3 | DSS | Spt6 | 937 | Spt6 | 773 |
| 3 | 3 | DSS | Spt6 | 956 | Spt6 | 319 |
| 1 | 3 | DSS | Spt6 | 956 | Spt6 | 1357 |
| 3 | 3 | DSS | Spt6 | 958 | Spt6 | 319 |
| 2 | 3 | DSS | Spt6 | 958 | Spt6 | 723 |
| 1 | 3 | DSS | Spt6 | 1007 | Spt6 | 319 |
| 3 | 3 | DSS | Spt6 | 1007 | Spt6 | 746 |
| 2 | 3 | DSS | Spt6 | 1007 | Spt6 | 749 |
| 3 | 3 | DSS | Spt6 | 1007 | Spt6 | 773 |
| 2 | 3 | DSS | Spt6 | 1047 | Spt6 | 1007 |
| 1 | 3 | DSS | Spt6 | 1130 | Spt6 | 728 |
| 3 | 3 | DSS | Spt6 | 1228 | Spt6 | 1239 |
| 1 | 3 | DSS | Spt6 | 1239 | Spt6 | 319 |
| 2 | 3 | DSS | Spt6 | 1239 | Spt6 | 611 |
| 3 | 3 | DSS | Spt6 | 1239 | Spt6 | 612 |
| 1 | 3 | DSS | Spt6 | 1274 | Spt6 | 1239 |
| 2 | 3 | DSS | Spt6 | 1299 | Spt6 | 1357 |
| 3 | 3 | DSS | Spt6 | 1355 | Spt6 | 1439 |
| 2 | 3 | DSS | Spt6 | 1355 | Spt6 | 1444 |
| 3 | 3 | DSS | Spt6 | 1366 | Spt6 | 1357 |
| 1 | 3 | DSS | Spt6 | 1378 | Spt6 | 1355 |

**Supplementary Table S5. Intra-protein crosslinks for Cdc73-10xHis used in the I-TASSER prediction of Cdc73 for structural modeling.**

| Crosslinker | AA 1 | AA 2 |
|---|---|---|
| EDC | 13 | 141 |
| EDC | 24 | 185 |
| EDC | 103 | 205 |
| EDC | 113 | 205 |
| EDC | 113 | 185 |
| EDC | 113 | 385 |
| EDC | 113 | 13 |
| EDC | 119 | 126 |
| EDC | 119 | 132 |
| EDC | 119 | 141 |
| EDC | 124 | 141 |
| EDC | 126 | 185 |
| EDC | 126 | 199 |
| EDC | 126 | 212 |
| EDC | 130 | 141 |
| EDC | 130 | 176 |
| EDC | 134 | 368 |
| EDC | 136 | 126 |
| EDC | 136 | 176 |
| EDC | 136 | 132 |
| EDC | 141 | 371 |
| EDC | 141 | 212 |
| EDC | 154 | 185 |
| EDC | 154 | 385 |
| EDC | 156 | 185 |
| EDC | 156 | 205 |
| EDC | 156 | 385 |
| EDC | 156 | 218 |
| EDC | 158 | 185 |
| EDC | 158 | 205 |
| EDC | 158 | 218 |
| EDC | 166 | 205 |
| EDC | 169 | 141 |
| EDC | 176 | 212 |
| EDC | 176 | 205 |

| | | |
|---|---|---|
| EDC | 185 | 176 |
| EDC | 185 | 141 |
| EDC | 185 | 126 |
| EDC | 185 | 134 |
| EDC | 194 | 141 |
| EDC | 197 | 205 |
| EDC | 199 | 176 |
| EDC | 205 | 141 |
| EDC | 205 | 134 |
| EDC | 218 | 141 |
| EDC | 218 | 132 |
| EDC | 226 | 141 |
| EDC | 236 | 261 |
| EDC | 261 | 236 |
| EDC | 263 | 24 |
| EDC | 263 | 16 |
| EDC | 348 | 366 |
| EDC | 371 | 132 |
| EDC | 385 | 134 |
| EDC | 385 | 126 |
| EDC | 385 | 129 |
| EDC | 385 | 132 |
| EDC | 385 | 141 |
| DSS | 21 | 13 |
| DSS | 77 | 17 |
| DSS | 119 | 130 |
| DSS | 119 | 136 |
| DSS | 119 | 185 |
| DSS | 119 | 13 |
| DSS | 119 | 205 |
| DSS | 119 | 304 |
| DSS | 124 | 185 |
| DSS | 124 | 13 |
| DSS | 124 | 304 |
| DSS | 124 | 385 |
| DSS | 124 | 371 |
| DSS | 130 | 263 |
| DSS | 130 | 185 |
| DSS | 130 | 205 |

| | | |
|---|---|---|
| DSS | 136 | 185 |
| DSS | 136 | 124 |
| DSS | 136 | 371 |
| DSS | 136 | 263 |
| DSS | 136 | 304 |
| DSS | 169 | 371 |
| DSS | 169 | 205 |
| DSS | 169 | 385 |
| DSS | 185 | 371 |
| DSS | 185 | 199 |
| DSS | 185 | 205 |
| DSS | 194 | 205 |
| DSS | 205 | 371 |
| DSS | 205 | 212 |
| DSS | 212 | 371 |
| DSS | 218 | 205 |
| DSS | 236 | 304 |
| DSS | 236 | 205 |
| DSS | 236 | 226 |
| DSS | 236 | 218 |
| DSS | 236 | 212 |
| DSS | 236 | 185 |
| DSS | 263 | 371 |
| DSS | 263 | 366 |
| DSS | 263 | 205 |
| DSS | 263 | 130 |
| DSS | 263 | 368 |
| DSS | 282 | 371 |
| DSS | 282 | 185 |
| DSS | 282 | 205 |
| DSS | 282 | 385 |
| DSS | 304 | 185 |
| DSS | 304 | 371 |
| DSS | 304 | 212 |
| DSS | 304 | 218 |
| DSS | 304 | 205 |
| DSS | 304 | 226 |
| DSS | 304 | 130 |
| DSS | 361 | 371 |

| | | |
|---|---|---|
| DSS | 366 | 371 |
| DSS | 366 | 17 |
| DSS | 368 | 366 |
| DSS | 371 | 366 |
| DSS | 371 | 212 |
| DSS | 371 | 205 |
| DSS | 371 | 368 |
| DSS | 371 | 13 |
| DSS | 385 | 371 |
| DSS | 385 | 185 |
| DSS | 385 | 212 |
| DSS | 385 | 205 |
| DSS | 385 | 17 |
| DSS | 385 | 130 |
| DSS | 385 | 218 |
| DSS | 385 | 366 |
| DSS | 385 | 368 |
| DSS | 385 | 263 |
